# Oxidation-Driven mtDNA B-Z Transition Activates ZBP1 to Mediate Acetaminophen Hepatotoxicity

**DOI:** 10.64898/2026.02.16.706077

**Authors:** Zhang-Hua Yang, Bo-Xin Zhang, Huan-Feng Ye, Rui Gong, Liang Shi, Zhi-Yu Cai, Qiang Chen, Lei Wu, Jia Huang, Le Zhang, Huipeng Jiao, Pinglong Xu, Qinjie Weng, Jie Zhang, Jinheng Pan, Shan Feng, Haibing Zhang, Xian Shen, Wei Mo

## Abstract

Acetaminophen (APAP) overdose induces mitochondrial damage in hepatocytes, leading to secondary toxicity that resists to N-acetylcysteine (NAC) treatment and culminates in hepatocyte death and acute liver failure (ALF)^1^. The underlying mechanisms remain poorly understood, often necessitating liver transplantation^2^. Here, we identify oxidative modification-driven B-to-Z transitions in mtDNA as the central pathological driver of APAP-induced ALF. Upon APAP exposure, oxidized mtDNA fragments leak into the cytosol, activating ZBP1 signaling via its Zα domain. Genetic inhibition of ZBP1 mitigates liver damage and improves survival. Using synthetic 12-bp dGdC duplexes, we demonstrate that 8-oxoG substitution, even under physiological salt concentrations, is sufficient to induce Z-DNA formation, enabling specific ZBP1 binding through its Zα domain. The 8-oxoG repair enzyme OGG1^3^, activated by TH10785^4^, removes 8-oxoG modifications and reverses Z-DNA to B-DNA conformation. In mice with lethal APAP toxicity, delayed NAC treatment results in 50% mortality. In contrast, TH10785 monotherapy increases survival to 90%, while its combination with NAC achieves 100% survival. These results define the oxidized mt Z-DNA-ZBP1 axis as a critical driver of APAP hepatotoxicity, providing fundamental insights into DNA conformational dynamics and therapeutic opportunities in drug-induced liver failure.

## Main

The liver serves as one of the principal detoxification organs in the human body, playing a critical role in the metabolism and elimination of toxins, drugs, and other deleterious substances. Nevertheless, the metabolic processing of drugs can also induce hepatic injury, a condition referred to as Drug-Induced Liver Injury (DILI)^5^. Acetaminophen (APAP), a widely utilized component in cold medications, can provoke extensive hepatocyte necrosis when consumed in excessive amounts. The central mechanism underlying its hepatotoxicity is mitochondrial dysfunction^6^. APAP is primarily metabolized by the cytochrome P450 enzyme CYP2E1, yielding the highly reactive and toxic metabolite N-acetyl-p-benzoquinone imine (NAPQI)^7^. NAPQI covalently binds to mitochondrial inner membrane proteins, disrupting the electron transport chain (ETC), altering mitochondrial membrane permeability, and triggering the overproduction of reactive oxygen species (ROS)^8,9^. These events culminate in mitochondrial impairment and subsequent hepatocyte necrosis. During the initial phase of APAP intoxication, the abundant availability of glutathione (GSH) facilitates the detoxification of NAPQI, thereby attenuating its direct deleterious effects on mitochondria^10^. However, beyond 8 hours post-APAP overdose, mitochondrial damage and ROS-driven oxidative stress initiate a cascade of extensive secondary injury to hepatocytes. This secondary damage cannot be mitigated by GSH administration beyond this critical 8-hour window, leading to irreversible and widespread hepatocyte death and the onset of acute liver failure (ALF)^1^. Therefore, elucidating the core pathological mechanisms underlying the secondary damage induced by APAP overdose is pivotal for developing strategies to reduce the incidence of APAP-induced acute liver failure.

### Cytoplasmic mtDNA activates ZBP1, not cGAS, to drive hepatocyte death

Mitochondrial damage-induced leakage of mitochondrial DNA (mtDNA) into the cytoplasm is a key trigger of immune responses and cell death, contributing to the pathogenesis of various diseases^11^. Similarly, APAP-induced mitochondrial injury leads to the cytosolic release of mtDNA. In both APAP-treated murine livers (**Extended Data Fig. 1a**) and the hepatocyte cell line AML12, a substantial accumulation of cytosolic double-stranded DNA (dsDNA) signals was detected (**Fig. 1a, b and Extended Data Fig. 1b, c**), which was confirmed to be mtDNA-derived (**Fig. 1a, b**).

**Fig. 1.**
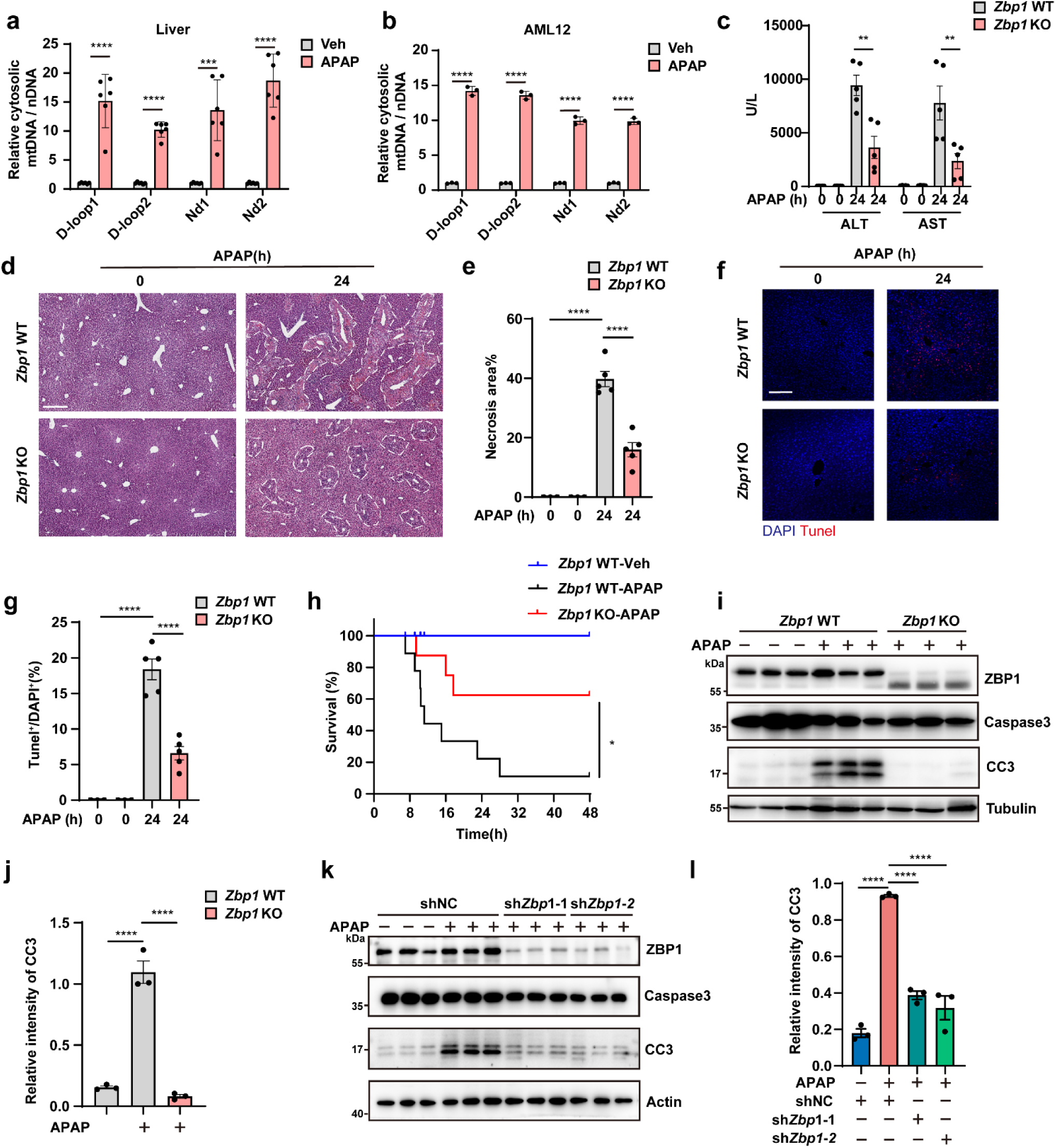
Cytoplasmic mtDNA activates ZBP1, not cGAS, to drive hepatocyte death. **a**,**b**, Relative amount of cytosolic mtDNA release in cytosol fraction of liver from WT mice (**a**) or AML12 cells (**b**) treated with or without APAP for 24 h. (**a**) 300 mg/kg i.p. for mice, n = 5 mice. (**b**) 10 mM for cell, n=3 independent samples. **c**, Serum ALT and AST levels were determined in *Zbp1* WT and *Zbp1* KO mice. 0 h: n=3 mice per genotype, 24 h: n=5 mice per genotype. **d**, Hematoxylin and eosin (H&E) staining of the liver sections from *Zbp1* WT and *Zbp1* KO mice. **e**, Necrotic areas were encircled and quantified. 0 h: n=3 mice, 24 h: n=5 mice. Scale bar, 300 μm. **f**,**g**, Representative images (**f**) and quantification analyses (**g**) of TUNEL staining in liver tissues from *Zbp1* WT and *Zbp1* KO mice. 0 h: n=3 mice, 24 h: n = 5 mice. Scale bar, 100 μm. **h**, Survival curves of *Zbp1* WT and *Zbp1* KO mice treated with a lethal dose of APAP (550 mg/kg, i.p.), n=8 mice per genotype. **i**,**j**, Western blotting (**i**) and quantification (**j**) analysis of liver proteins from *Zbp1* WT and *Zbp1* KO mice treated with or without APAP for 24 h. n=3 mice per group. CC3: cleaved caspase-3. **k**,**l**, Western blotting (**k**) and quantification (**l**) analysis of shNC (non-target control) or sh*Zbp1* AML12 cells treated with or without APAP for 24 h. n=3 independent samples. Data are mean ± s.e.m. Statistical analysis was performed using one-way analysis of variance (ANOVA) (**j** and **l**), two-way analysis of variance (ANOVA) (**a**, **b**, **c**, **e** and **g**), and Mantel−Cox tests (**h**); ns, not significant; \**P* < 0.05; \*\**P* < 0.01; \*\*\**P* < 0.001; \*\*\*\**P* < 0.0001.

Under normal conditions, aberrantly leaked cytosolic mtDNA is recognized by cyclic GMP-AMP synthase (cGAS), triggering innate immune signaling^12^. However, in APAP-treated cells, cytosolic mtDNA failed to activate cGAS, as indicated by the absence of increased cGAMP production (**Extended Data Fig. 1d, e**)—the enzymatic product of cGAS activation—and the lack of phosphorylation of its downstream effector, TANK-binding kinase 1 (TBK1) (**Extended Data Fig. 1f**). Consistent with these findings, genetic ablation of cGAS did not confer protection against APAP-induced liver injury in mice. Compared to wild-type (WT) controls, cGAS-deficient mice showed no improvement in serum markers of liver function, such as alanine aminotransferase (ALT) and aspartate aminotransferase (AST) (**Extended Data Fig. 1g**), nor a reduction in TUNEL-positive hepatocytes or necrotic areas (**Extended Data Fig. 1h-k**). These findings may be attributed to the extremely low expression levels of key components of DNA-sensing pathways in hepatocytes^13,14^, particularly the cGAS, as previously reported and as observed in our study.

Other cytosolic DNA sensors, including interferon gamma-inducible protein 16 (IFI16), DDX41, and absent in melanoma 2 (AIM2), are known to activate inflammatory cytokine responses^15–17^. However, 24 hours post-APAP treatment, no significant changes were observed in the levels of interferons or pro-inflammatory cytokines such as tumor necrosis factor (TNF), interleukin-1β (IL-1β), and interleukin-6 (IL-6) (**Extended Data Fig. 1l**). These findings suggest that APAP-induced cytosolic mtDNA does not trigger conventional innate immune signaling pathways.

Intriguingly, Z-DNA binding protein 1 (ZBP1), a non-canonical cytosolic dsDNA sensor, does not primarily induce type I interferon responses but is instead implicated in the direct initiation of cell death^18,19^. Based on these observations, we hypothesized that cytosolic mtDNA released upon APAP treatment is sensed by ZBP1, which subsequently drives hepatocyte death.

In the APAP-induced liver injury model, genetic ablation of *Zbp1* confers significant protection against hepatic dysfunction in mice as indicated by ALT and AST (**Fig. 1c**). Histopathological analysis using hematoxylin and eosin (H&E) staining demonstrated a substantial reduction in necrotic areas in the livers of *Zbp1* knockout (KO) mice compared to those of WT controls (**Fig. 1d, e**). Quantification of TUNEL staining further revealed that the proportion of TUNEL-positive cells in *Zbp1* KO livers decreased to 5%, indicating a marked reduction in cell death (**Fig. 1f, g**). Additionally, following APAP administration, the infiltration of immune cells, particularly neutrophils, was significantly attenuated in the livers of *Zbp1* KO mice relative to WT mice (**Extended Data Fig. 1m, n**). In a survival study, treatment with a lethal dose of APAP resulted in a 90% mortality rate within 48 hours in WT mice; however, *Zbp1* KO reduced the mortality rate to 40% (**Fig. 1h**), highlighting the protective role of ZBP1 deficiency in mitigating APAP-induced lethality.

Building on these findings, we further elucidated the mechanism through which ZBP1 drives hepatocyte cell death. Although ZBP1 is known to recruit RIPK3 via its RHIM domain to initiate necroptosis^20^, this pathway is not operational in hepatocytes due to the absence of RIPK3 expression, and accordingly, necroptosis (pMLKL) was not observed in hepatocytes following APAP treatment (**Extended Data Fig. 2a**). Consistent with this, genetic ablation of RIPK3 or MLKL failed to confer protection against APAP-induced liver injury (**Extended Data Fig. 2b**), ruling out necroptosis as a contributing mechanism. Beyond necroptosis, ZBP1 has been implicated in the regulation of apoptosis. In APAP-treated livers, robust caspase-3 cleavage was detected, whereas *Zbp1* KO livers exhibited no such activation of caspase-3 (**Fig. 1i, j**), suggesting a critical role for ZBP1 in APAP-induced apoptotic cell death. To address the potential confounding effects of non-hepatocyte contributions in liver tissue extracts, we validated these findings in a hepatocyte cell line, where APAP treatment similarly induced caspase-3 cleavage, and *Zbp1* knockdown effectively suppressed apoptosis (**Fig. 1k, l**). We also explored the possibility of pyroptosis in this context. However, no evidence of Gasdermin D-mediated pyroptosis was detected in APAP-treated livers (**Extended Data Fig. 2c**), further corroborating the lack of AIM2 activation despite the presence of cytosolic mtDNA. Additionally, Gasdermin E, whose cleavage is caspase-3-dependent, remained intact following APAP treatment (**Extended Data Fig. 2b**), reinforcing the notion that APAP-induced caspase-3 activation specifically drives apoptosis rather than Gasdermin E-mediated pyroptosis. Collectively, these findings indicate that the critical pathological mechanism underlying APAP-induced liver injury involves the release of mtDNA into the cytoplasm, which subsequently activates the ZBP1 pathway, leading to hepatocyte death.

### ZBP1 binds to Z-DNA derived from released mitochondrial DNA

ZBP1 functions as a critical sensor of left-handed Z-form nucleic acids (Z-DNA/Z-RNA)^21–24^, raising the possibility that APAP-induced hepatocyte injury triggers the formation of Z-nucleic acids (Z-NA). To investigate this hypothesis, we employed the Z22 antibody, which selectively recognizes Z-NA, including both Z-DNA and Z-RNA, to detect their presence in APAP-treated hepatocytes. As shown in **Fig. 2a-c**, significant accumulation of Z-NA was observed in both liver tissues, ex vivo hepatocyte and cultured hepatocyte AML12 cells following APAP exposure. Notably, the Z-NA signal detected in the livers of APAP-treated mice, primary hepatocyte and APAP-stimulated AML12 cells was abolished by DNase I treatment but remained unaffected by RNase A (**Fig. 2a-f**), confirming that the observed Z-NA signal was primarily derived from DNA. The APAP-induced Z-DNA accumulation was comparable between WT and *Zbp1* KO mice (**Extended Data Fig. 3a, b**), indicating that Z-DNA formation occurs independently of ZBP1.

**Fig. 2.**
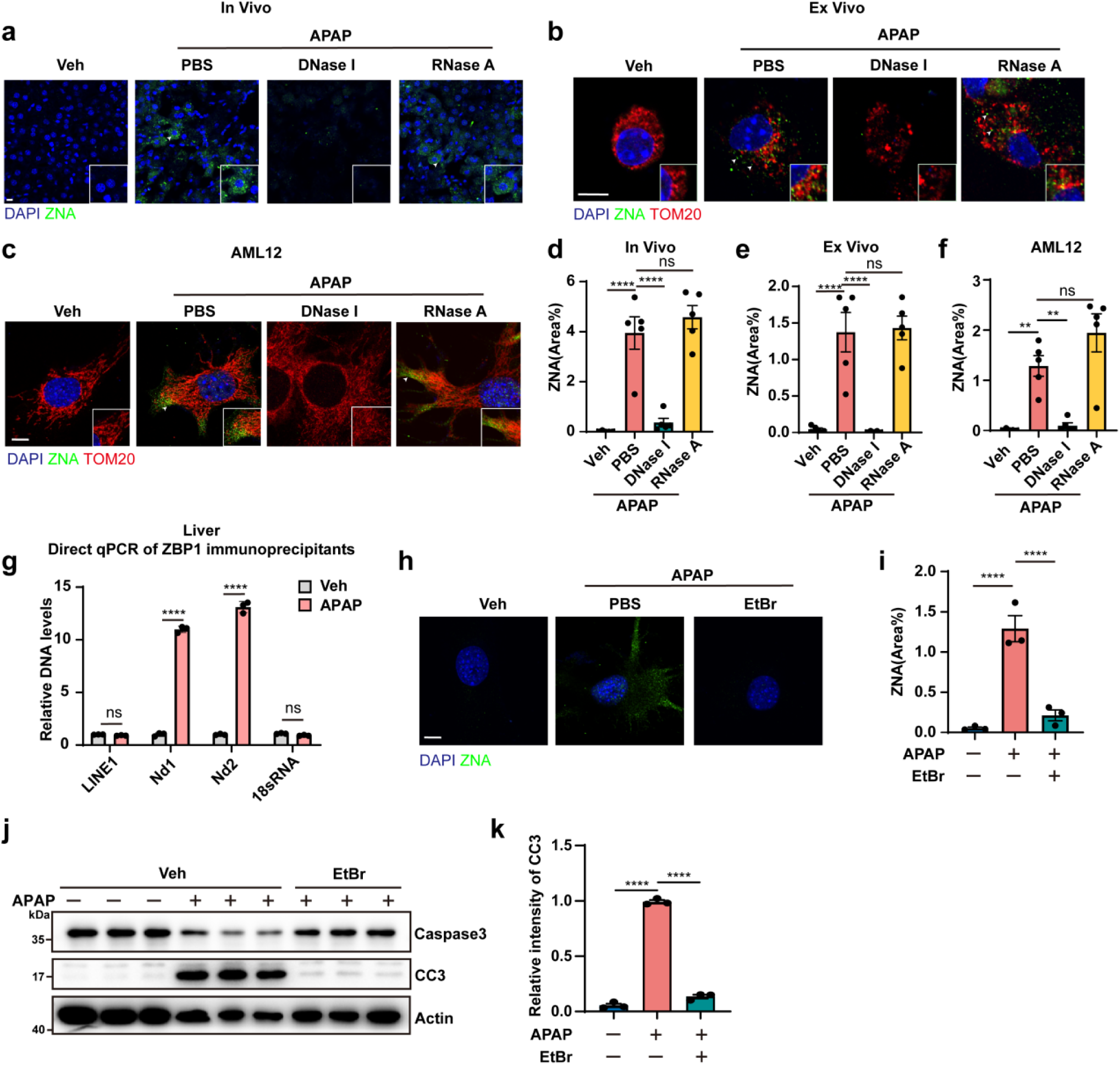
ZBP1 binds to Z-DNA derived from released mitochondrial DNA. **a**, Immunofluorescence staining of Z-NA (green) with anti-Z-NA antibody with or without DNase I (25 U/mL) or RNase A (2 mg/mL) treatment in the liver of WT mice treated with Vehicle or APAP (300 mg/kg) for 24 h. Scale bar, 10 μm. **b**, Immunofluorescence staining of Z-NA (green) and TOM20 (red) in the hepatocytes isolated from WT mice treated with Vehicle or APAP (300 mg/kg) for 24 h. Scale bar, 10 μm. **c**, Immunofluorescence staining of Z-NA (green) and TOM20 (red) in the AML12 cells treated with Vehicle or APAP (10 mM) for 24 h. Scale bar, 10 μm. a-c, The white arrow indicates the released mitochondrial DNA (mtDNA). **d**, Quantification of the fluorescence area of Z-NA signals in (**a**). n=5 mice per group. **e**, Quantification of the fluorescence area of Z-NA signals in (**b**). n=5 mice per group. **f**, Quantification of the fluorescence area of Z-NA signals in (**c**). n=5 independent samples. **g**, Quantitative qPCR analysis of ZBP1-binding DNAs enriched by anti-ZBP1 immunoprecipitation in liver of WT mice treated with Vehicle or APAP (300 mg/kg) for 24 h. n=3 mice per group. **h**, Immunofluorescence staining of Z-NA (green) in the AML12 cells with or without mtDNA depletion by EtBr (400 ng/ml) treatment for 6 days and stimulated with Vehicle or APAP (10 mM) for 24 h. Scale bar, 10 μm. **l**, Quantification of the fluorescence area of Z-NA signals in (**h**). n=3 independent samples. **j**,**k**, Western blotting (**j**) and quantification (**k**) analysis of the AML12 cells with or without mtDNA depletion by EtBr (400 ng/ml) treatment for 6 days and stimulated with Vehicle or APAP (10 mM) for 24 h. n=3 independent samples. Data are mean ± s.e.m. Statistical analysis was performed using one-way analysis of variance (ANOVA) (**d**, **e**, **f**, **I** and **k**) and two-way analysis of variance (ANOVA) (**g)**; ns, not significant; \**P* < 0.05; \*\**P* < 0.01; \*\*\**P* < 0.001; \*\*\*\**P* < 0.0001.

A substantial fraction of Z-DNA colocalized with the mitochondrial marker TOM20, while a portion was detected outside mitochondria (**Fig. 2b, c**), suggesting that the Z-DNA may originate from leaked mtDNA. To directly determine the source of Z-DNA that engages ZBP1 upon APAP stimulation, we performed ZBP1 immunoprecipitation (IP) followed by qPCR analysis to distinguish nuclear versus mitochondrial DNA (**Fig. 2g** and **Extended Data Fig. 3c**). The results demonstrated that, following APAP exposure, ZBP1 was specifically bound to mtDNA, as indicated by the enrichment of mitochondrial markers Nd1 and Nd2, while markers of nuclear DNA, including LINE1 and RNA18S, were absent (**Fig. 2g**). Consistent with this, depletion of mtDNA using ethidium bromide (EB) (**Extended Data Fig. 3d**) abolished APAP-induced Z-DNA formation (**Fig. 2h, i**). Furthermore, EB-mediated mtDNA depletion significantly attenuated APAP-induced apoptotic signaling (**Fig. 2j, k**), reinforcing the critical role of mtDNA-derived Z-DNA in promoting cell death pathways.

The N-terminal Zα domain of ZBP1 is essential for its interaction with double-stranded DNA (dsDNA), and point mutations in the Zα1 and Zα2 subdomains effectively disrupted this binding. In the APAP-induced liver injury model, mice expressing ZBP1 Zα mutants exhibited hepatoprotection comparable to that observed in *Zbp1* KO mice (**Extended Data Fig. 4a**). This protection was characterized by reduced apoptosis (**Extended Data Fig. 4b-e**), diminished necrotic regions (**Extended Data Fig. 4f, g**), decreased immune cell infiltration (**Extended Data Fig. 4h, i**). These findings indicate that ZBP1-mediated recognition of Z-DNA plays a pivotal role in APAP-induced hepatocyte death and liver injury, underscoring the pathogenic significance of mtDNA-derived Z-DNA in APAP toxicity.

### Oxidative stress-induced mtDNA fragmentation and cytoplasmic leakage in APAP-induced liver injury

In the context of APAP-induced oxidative stress, excessive reactive oxygen species (ROS) mediate mtDNA oxidation, leading to its fragmentation (**Fig. 3a**). 8-Oxoguanine (8-hydroxyguanine, 8-oxoG) is one of the most common DNA lesions resulting from ROS. Using an 8-oxoG-specific antibody, we observed a significant elevation in cytoplasmic oxidized DNA levels in the APAP-induced liver injury model (**Fig. 3b**). Additionally, APAP-treated hepatocytes exhibited an approximately 16-fold increase in cytoplasmic oxidized DNA (**Fig. 3c**). Notably, treatment with the mitochondrial-targeted ROS scavenger MitoQ markedly reduced cytoplasmic oxidized DNA levels (**Fig. 3d-f**). This reduction was accompanied by a substantial decrease in both intra-mitochondrial and cytoplasmic Z-DNA (**Fig. 3g, h**), suggesting that mtDNA oxidation promotes Z-DNA formation. Consistently, ROS depletion by MitoQ also significantly attenuated APAP-induced apoptotic signaling (**Fig. 3i, j**).

**Fig. 3.**
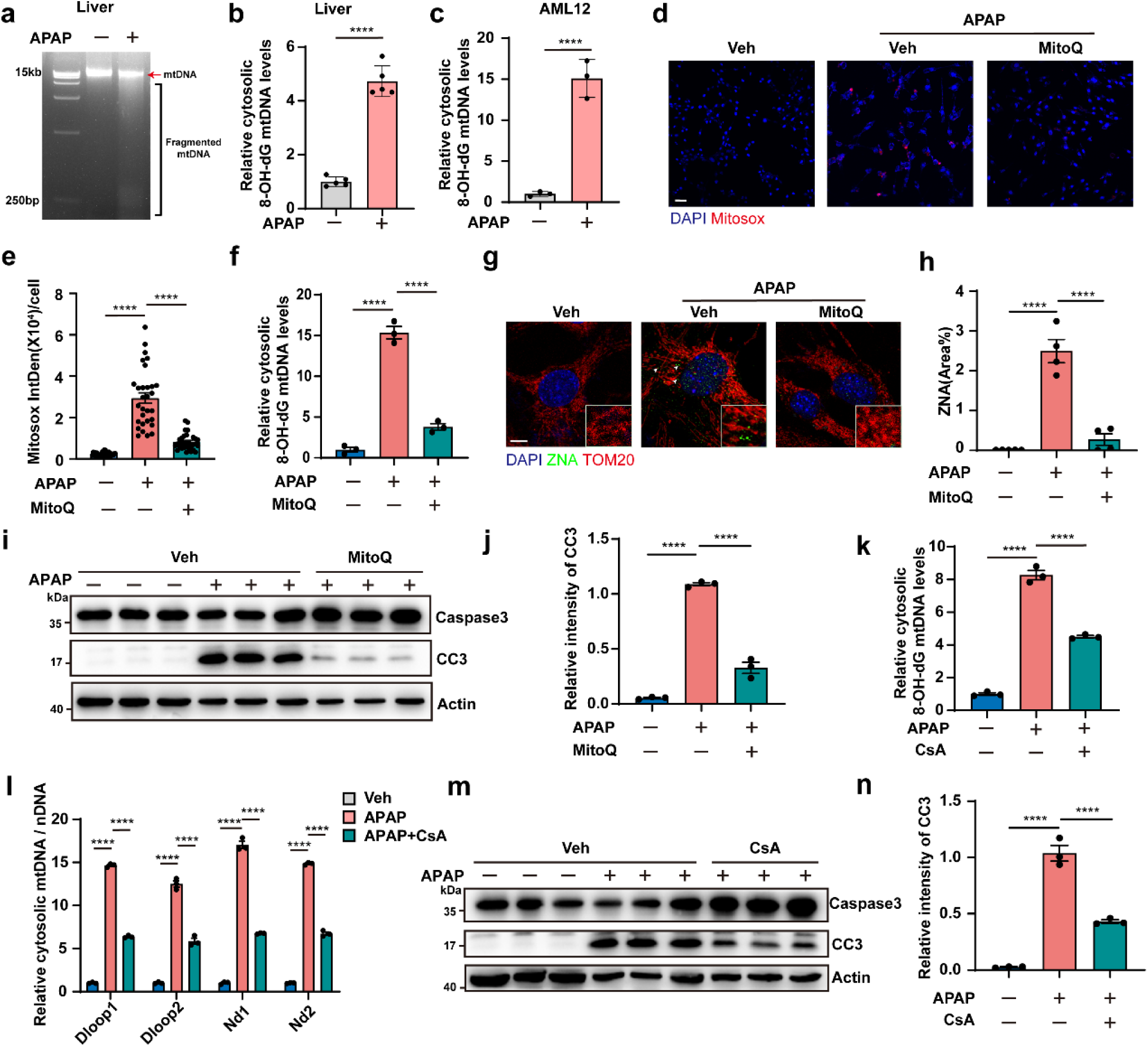
Oxidative stress-induced mtDNA fragmentation and cytoplasmic leakage in APAP-induced liver injury. **a**, Agarose gel electrophoresis of cytosolic mtDNA from liver of WT mic treated with or without APAP (300 mg/kg) for 24 h. **b**, Quantification of the amounts of cytosolic Ox-mtDNA in the liver of WT mic treated with or without APAP (300 mg/kg) for 24 h. n=5 mice per group. **c**, Quantification of the amounts of cytosolic Ox-mtDNA in the AML12 cells treated with or without APAP (10 mM) for 24 h. n=3 independent samples. **d**, Representative images of mitoSOX Red+ cells indicating mitochondrial ROS production in AML12 cells treated with or without APAP (10 mM) for 24 h, in the presence or absence of MitoQ (1 μM). Scale bars, 50 μm. **e**, Quantification analysis of the fluorescence intensity of MitoSOX Red per cell in (**d**). n=10 cells per group, repeated 3 times independently with similar results. **f**, Quantification of the amounts of cytosolic Ox-mtDNA in the AML12 cells treated with or without APAP (10 mM) for 24 h, in the presence or absence of MitoQ (1 μM). n=3 independent samples. **g**, Immunofluorescence staining of Z-NA (green) and TOM20 (red) in the AML12 cells treated with or without APAP (10 mM) for 24 h, in the presence or absence of MitoQ (1 μM). The white arrow indicates the released mtDNA. **h**, Quantification of the fluorescence area of Z-NA signals in (**g**). n=4 independent samples. **i**,**j**, Western blotting (**i**) and quantification (**j**) analysis of the AML12 cells treated with or without APAP (10 mM) for 24 h, in the presence or absence of MitoQ (1 μM). n=3 independent samples. **k**, Quantification of the amounts of cytosolic Ox-mtDNA in the AML12 cells treated with or without APAP (10 mM) for 24 h, in the presence or absence of CsA (10 μM). n=3 independent samples. **i**, Quantitative PCR analysis of cytosolic mtDNA release in cytosol fraction of AML12 cells treated with or without APAP (10 mM) for 24 h, in the presence or absence of CsA (10 μM). n=3 independent samples. **m**,**n**, Western blotting (**m**) and quantification (**n**) analysis of the AML12 cells treated with or without APAP (10 mM) for 24 h, in the presence or absence of CsA (10 μM). n=3 independent samples. Data are mean ± s.e.m. Statistical analysis was performed using Unpaired two-tailed Student’s *t*-test (**b, c**), two-way analysis of variance (ANOVA) (**l**), and one-way analysis of variance (ANOVA) (**e, f, h, j, k** and **n**); ns, not significant; \**P* < 0.05; \*\**P* < 0.01; \*\*\**P* < 0.001; \*\*\*\**P* < 0.0001.

Next, we sought to elucidate the mechanism underlying oxidized mtDNA release. Previous studies have proposed that oxidized mtDNA undergoes cleavage by Flap Endonuclease 1 (FEN1) to generate short oxidized mtDNA fragments before their release into the cytoplasm^25^. However, in APAP-treated hepatocytes, downregulation of FEN1 did not affect either the cytoplasmic leakage of oxidized mtDNA (**Extended Data Fig. 5a, b**) or the extent of APAP-induced hepatocyte apoptosis (**Extended Data Fig. 5c, d**). These findings suggest that mtDNA fragmentation in APAP-treated cells occurs independently of FEN1-mediated processing.

Additionally, it has been suggested that short oxidized mtDNA fragments are transported into the cytoplasm via the mitochondrial permeability transition pore (mPTP)-VDAC1 transport axis^26^. In our study, pharmacological inhibition of mPTP using cyclosporine A (CsA) resulted in a partial reduction in cytoplasmic oxidized mtDNA levels (**Fig. 3k, l**) and a corresponding attenuation of APAP-induced apoptosis (**Fig. 3m, n**). While previous reports indicate that mtDNA promotes VDAC1 oligomerization, thereby forming ion channels that facilitate mtDNA leakage from the mitochondrial intermembrane space to the cytoplasm^26^, treatment with the VDAC1 oligomerization inhibitor VBIT-4 failed to reduce cytoplasmic oxidized mtDNA levels or mitigate APAP-induced apoptosis (**Extended Data Fig. 5e-g**).

These findings suggest that APAP-induced mtDNA fragmentation and cytoplasmic leakage occur through a distinct mechanism. We hypothesize that severe oxidative stress triggered by APAP leads to extensive mtDNA oxidation, promoting fragmentation in a nuclease-independent manner. Furthermore, the resulting oxidized mtDNA fragments may translocate into the cytoplasm via a pathway that is not fully dependent on the mPTP-VDAC1 transport system.

### Oxidative modifications of DNA bases facilitate the B-to-Z DNA conformational transition

The administration of MitoQ for ROS clearance resulted in a reduction in Z-DNA levels, establishing a direct link between oxidative stress and the B-to-Z conformational transition of DNA. In AML12 cells, following APAP treatment, Z-DNA and 8-oxoG co-localized within both mitochondria and the cytoplasm (**Fig. 4a, b**), supporting the notion that oxidative modifications drive the B-to-Z DNA transition.

**Fig. 4.**
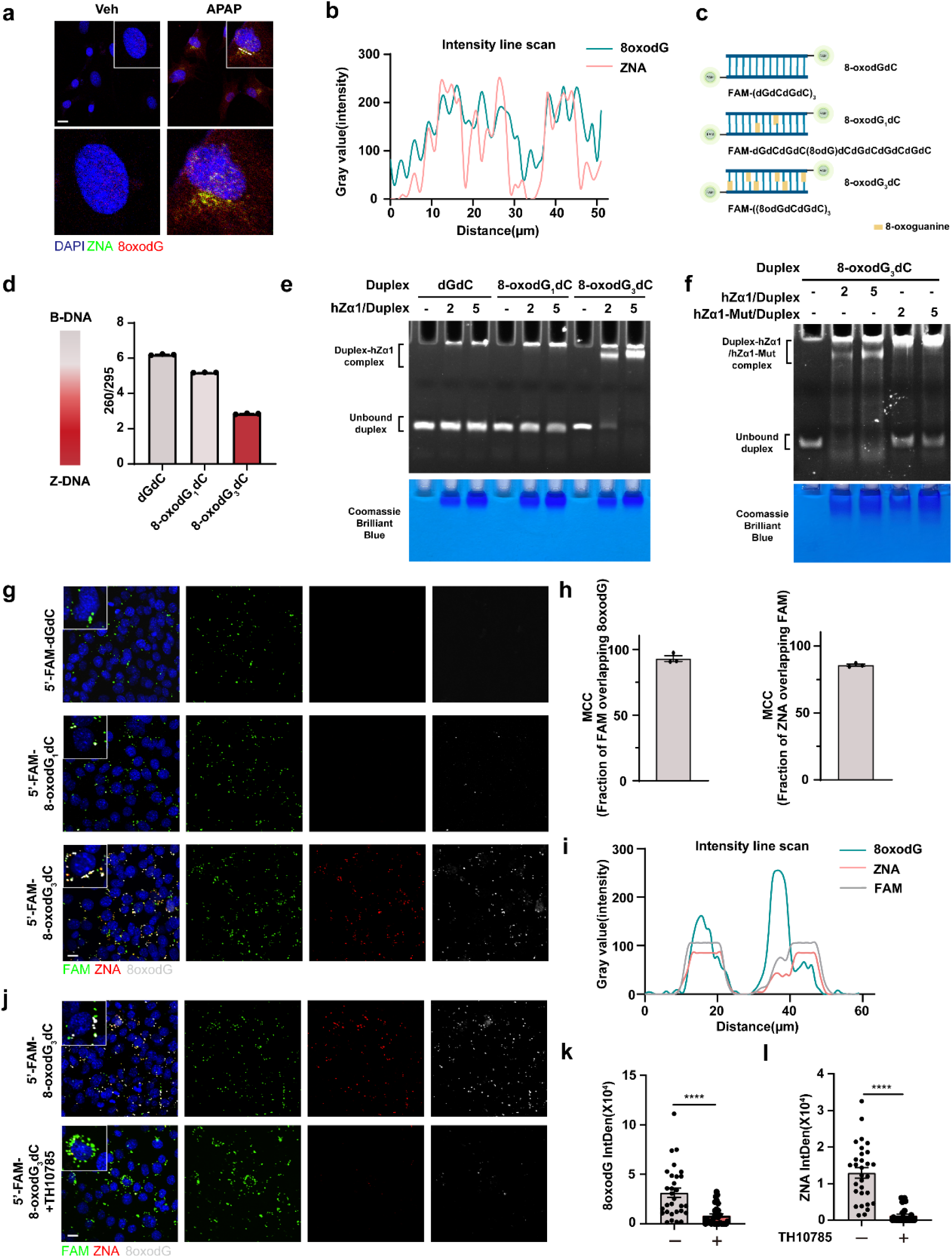
Oxidative modifications of DNA bases facilitates the B-to-Z DNA conformational transition. **a**, Immunofluorescence staining of Z-NA (green) and 8oxodG (red) in the AML12 cells treated with or without APAP (10 mM) for 24 h. Dotted line indicates region chosen for intensity line scan analysis for (**b**). Scale bars, 20 μm. **b**, Normalized intensity line scan of ZNA and 8oxodG in (**a**). **c**, Schematics of the synthesized dsDNA sequences (12 bp) containing six consecutive GC repeats with defined oxidation levels (0, 1, or 3 8-oxoG substitutions per strand). **d**, Change of A260/295 as a measure of B-DNA (high ratio ∼ 6) versus Z-DNA (lower ratio of ∼3) formation is graphed. **e**,**f,** EMSA assays of dGdC, 8-oxodG_1_dC, and 8-oxodG_3_dC duplexes incubated with (**e**) 1:2,1:5 molar ratio of duplex to Zα1 domain from human ZBP1 (hZα1), and (**f**) 8-oxodG_3_dC duplexes incubated with 1:2,1:5 molar ratio of duplex to hZα1 and hZα1-Mut. [Oligonucleotide] = 1 μM. **g**, Immunofluorescence staining of FAM (green), ZNA (red) and 8oxodG (gray) in the *Zbp1* KO MEFs transfected with dGdC, 8-oxodG_1_dC and 8-oxodG_3_dC duplexes respectively. Dotted line indicates region chosen for intensity line scan analysis for (**l**). **h**, Quantification of percent of cells with FAM and 8oxodG or ZNA overlap signal in (**g**). n=3 independent experiments. **i**, Normalized intensity line scan of FAM, ZNA and 8oxodG in (**g**). **j**, Immunofluorescence staining of FAM (green), ZNA (red) and 8oxodG (gray) in the *Zbp1* KO MEFs transfected with 8-oxodG_3_dC duplexes for 24 h in the presence or absence of TH10785 (10 μM). **k,l**, Quantification of the fluorescence intensity of 8oxodG (**k**) and ZNA (**l**) in (**j**). n=10 cells per group, repeated 3 times independently with similar results. Data are mean ± s.e.m. Statistical analysis was performed using Unpaired two-tailed Student’s *t*-test (**k** and **l**); ns, not significant; \**P* < 0.05; \*\**P* < 0.01; \*\*\**P* < 0.001; \*\*\*\**P* < 0.0001.

To determine whether oxidative modifications alone are sufficient to induce the B-to-Z transition, we synthesized short double-stranded DNA (dsDNA) sequences (12 bp) containing six consecutive GC repeats, incorporating defined oxidation levels (0, 1, or 3 8-oxoG substitutions per strand) (**Fig. 4c**). The Z-DNA conformation was assessed using the A260/295 absorbance ratio as a readout, which is markedly reduced in Z-DNA relative to the canonical B-form. The unmodified dsDNA initially displayed an A260/295 ratio of approximately 6, consistent with the B-form conformation (**Fig. 4d**). Upon substitution of guanine (G) with 8-oxoG, an inverse relationship emerged between the oxidation level and the A260/295 ratio, signifying a progressive transition from the B-form to the Z-conformation (**Fig. 4d**). Notably, while the canonical B-to-Z transition typically requires high-salt conditions (≥2 M NaCl), this oxidation-driven structural shift occurred at low-salt concentrations (20 mM NaCl), underscoring the unique ability of oxidative modifications to induce Z-DNA formation under physiologically relevant conditions (**Fig. 4d**). The B-to-Z transition induced by oxidative modification was further validated through electrophoretic mobility shift assays (EMSA), which revealed specific binding of 8-oxoG-modified DNA to ZBP1 (**Fig. 4e**). Notably, this interaction was strictly dependent on the Zα domain of ZBP1, as evidenced by experiments using a mutated Za domain that had lost its ability to bind Z-DNA (**Fig. 4f**). This domain-specific binding provides definitive evidence that the oxidized DNA adopts a left-handed Z-conformation, rather than merely representing a chemically modified B-form structure. Of note, the efficiency of Z-DNA formation was markedly enhanced in DNA strand containing three 8-oxoG substitutions compared to those with a single substitution (**Fig. 4e**), further underscoring the critical role of cumulative oxidative modifications in driving the B-to-Z transition.

To further determine whether oxidative modification of DNA alone is sufficient to induce Z-DNA formation in cells, we transfected oxidized dsDNA into *Zbp1* KO mouse embryonic fibroblasts (MEFs). These cells lack both known Zα-binding proteins, ZBP1 and ADAR1-p150, with the latter exhibiting minimal expression in basal conditions^27^. In alignment with our biochemical data, the oxidized dsDNA adopted the Z-conformation intracellularly in the absence of ZBP1 and ADAR1-p150 (**Fig. 4g-i**). Thus, Z-DNA formation by oxidized dsDNA occurred independently of the stabilization by ZNA binding protein. Consistent with the in vitro results (**Fig. 4e**), Z-DNA formation in cells was significantly more efficient in DNA strands containing three 8-oxoG substitutions compared to those with a single substitution (**Fig. 4g**), further underscoring the ability of oxidative modifications to promote the B-to-Z transition in a cellular context. Together, these findings robustly demonstrate that oxidative modifications alone are sufficient to drive the B-to-Z transition, both in vitro and in cells.

To investigate the reversibility of oxidative modifications in B-to-Z DNA transitions, we employed TH10785, a recently identified agonist of OGG1, to activate this DNA glycosylase that specifically removes oxidized guanine residues^4^. In *Zbp1* KO MEF cells transfected with oxidized dsDNA, OGG1 activation led to efficient removal of oxidative modifications, as evidenced by the loss of 8-oxoG signals (**Fig. 4j, k**). Concomitantly, the DNA reverted from the Z-form to the B-form, as indicated by the disappearance of Z-DNA signals (**Fig. 4j, l**). These findings provide compelling evidence that oxidative modifications not only sufficiently drive the B-to-Z DNA transition but also that this structural shift can be reversed through enzymatic deoxidation by OGG1.

### Deoxidation facilitates Z-to-B DNA transition, protecting against APAP-induced liver failure and mortality

To assess the pathological relevance of oxidation-driven B-to-Z DNA transitions in APAP-induced liver injury, we examined hepatocyte cell lines following APAP treatment. APAP exposure promoted the formation of oxidized Z-DNA, while OGG1 agonist administration efficiently reversed these oxidative modifications, restoring DNA to its B-form conformation (**Fig. 5a-c**). This intervention significantly reduced cytoplasmic oxidized mtDNA levels and diminished Z-DNA content in both mitochondrial and cytoplasmic fractions (**Fig. 5a**). Notably, these molecular resulted in substantially attenuated cellular apoptosis (**Extended Data Fig. 6a, b**).

**Fig. 5.**
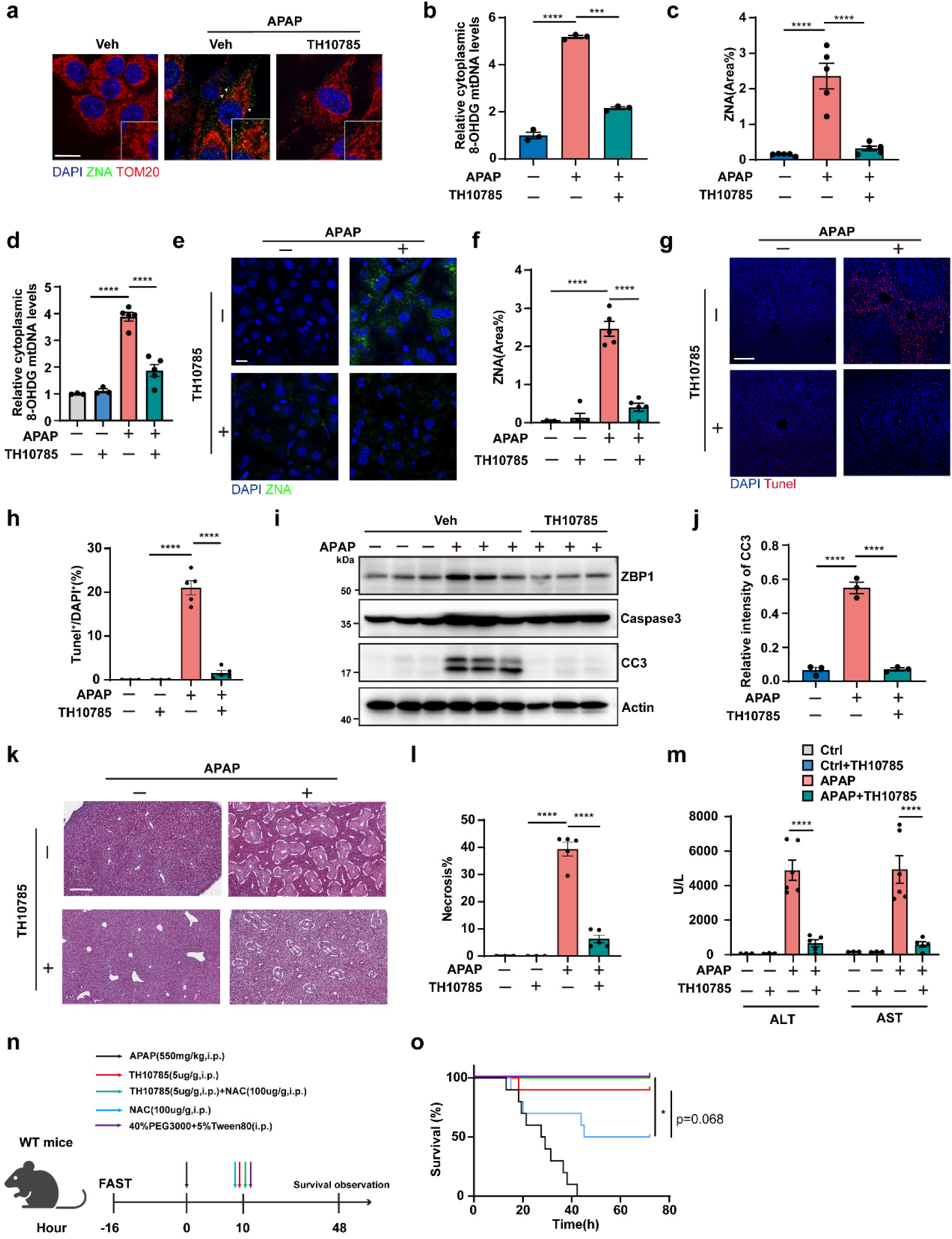
Deoxidation facilitates Z-to-B DNA transition, protecting against APAP-induced liver failure and mortality. **a**, Immunofluorescence staining of Z-NA (green) and TOM20 (red) in AML12 cells treated with or without APAP (10 mM) for 24 h in the presence or absence of TH10785 (10 μM). The white arrow indicates the released mtDNA. **b**, Quantification of the amounts of cytosolic Ox-mtDNA in AML12 cells treated with or without APAP (10 mM) for 24 h, in the presence or absence of TH10785 (10 μM). n=3 independent samples. **c**, Quantification of the fluorescence area of Z-NA signals in (**a**). n=5 independent samples. **d**, Quantification of the amounts of cytosolic Ox-mtDNA in the liver of WT mice treated with or without APAP (300 mg/kg) in the presence or absence of TH10785 (5 μg/g) for 24 h. n=5 mice per group. **e**,**f**, Immunofluorescence staining (**e**) and quantification (**f**) of Z-NA (green) in the liver of WT mice treated with or without APAP (300 mg/kg) for 24 h in the presence or absence of TH10785 (5 μg/g). Scale bars, 15 μm. n=5 mice per group. **g**,**h** Representative images (**g**) and quantification analyses (**h**) of TUNEL staining in liver tissues from WT mice treated with or without APAP (300 mg/kg) in the presence or absence of TH10785 (5 μg/g) for 24 h. Scale bar, 100 μm. **i**,**j**, Western blotting (**i**) and quantification (**j**) analysis of liver proteins from WT mice treated with or without APAP (300 mg/kg) in the presence or absence of TH10785 (5 μg/g) for 24 h. n=3 mice per group. **k**, Hematoxylin and eosin (H&E) staining of the liver sections from WT mice treated with or without APAP (300 mg/kg) for 24 h in the presence or absence of TH10785 (5 μg/g). **l**, Necrotic areas were encircled and quantified. Scale bar, 300 μm. **m,** Serum ALT and AST levels were determined in WT mice treated with or without APAP (300 mg/kg) for 24 h in the presence or absence of TH10785 (5 μg/g). **d-m**, WT mice only treated with or without TH10785: n=3; WT mice treated with or without APAP in the presence or absence of TH10785: n=5. **n**, The scheme of APAP treatment in the presence or absence of TH10785 or NAC. **o**, Survival curves of WT mice treated with a lethal dose of APAP (550 mg/kg i.p.) in the presence or absence of TH10785 or NAC, n=10 mice per group. Data are mean ± s.e.m. Statistical analysis was performed using one-way analysis of variance (ANOVA (**b, c, d, h, j,** and **l**), two-way analysis of variance (ANOVA) (**m**), and Mantel−Cox tests (**o**); ns, not significant; \**P* < 0.05; \*\**P* < 0.01; \*\*\**P* < 0.001; \*\*\*\**P* < 0.0001.

Consistently, in an APAP-induced liver injury model, treatment with the OGG1 agonist substantially decreased cytoplasmic oxidized mtDNA levels (**Fig. 5d**), which was accompanied by a corresponding reduction in Z-DNA (**Fig. 5e, f**). This molecular effect was associated with reduced apoptosis (as indicated by TUNEL staining and cleaved caspase-3) (**Fig. 5g-j**), diminished necrotic regions (**Fig. 5k, l**), decreased immune cell infiltration (**Extended Data Fig. 6c-f**), and overall preservation of liver function (**Fig. 5m**). Notably, the OGG1 agonist did not affect APAP metabolism in cells or the liver, as indicated by CYP2E1 (the key APAP-metabolizing enzyme) levels and GSH levels (**Extended Data Fig. 7a-d**). Additionally, it did not influence ROS production (**Extended Data Fig. 7e, f**) or ZBP1 expression levels (**Extended Data Fig. 7g**), confirming that its protective effects were specifically mediated through the direct removal of oxidative modifications from guanine in DNA.

To evaluate the therapeutic potential of the OGG1 agonist in APAP-induced liver injury, we first examined its efficacy at a lethal APAP dose. Treatment with the OGG1 agonist conferred 90% protection against mortality (**Extended Data Fig. 8a, b**). To simulate a clinically relevant scenario, we employed a delayed-treatment model in which intervention was initiated 10 hours post-APAP administration (**Fig. 5n**). In this setting, the OGG1 agonist maintained a 90% survival rate, significantly surpassing the efficacy of N-acetylcysteine (NAC), the current clinical standard, which provided less than 50% protection (**Fig. 5o**). Strikingly, combining the OGG1 agonist with NAC resulted in 100% survival (**Fig. 5o**), demonstrating a synergistic therapeutic effect and underscoring the potential of this dual approach for clinical translation.

These findings highlight the pivotal role of oxidative modifications in APAP-induced liver injury and establish that promoting deoxidation via OGG1 activation facilitates the Z-to-B DNA transition, effectively mitigating hepatotoxicity and reducing mortality. This strategy represents a promising therapeutic avenue for improving clinical outcomes in APAP-induced liver failure.

## Acknowledgements

We thank professor Lijian Hui (Chinese Academy of Sciences) for kindly provided us with proliferating human hepatocytes (ProliHHs). The study was supported by the National Natural Science Foundation of China (82588302, 32225016, 82530046 to W.M.; 32170751, 92581110 to Z.-H.Y.; 82203426, 82472742 to L.S.), the National Key R&D Program of China (2024YFA1306400 and 2021YFA1101401 to W.M.), the Zhejiang Provincial Natural Science Foundation of China (LHZSD25C070001 to W.M.), the “Pioneer” and “Leading Goose” R&D Program of Zhejiang (No.2025C02110 to W.M.), the Noncommunicable Chronic Diseases-National Science and Technology Major Project (2024ZD0524900 to Z.-H.Y.).

## Author contributions

Z.-H.Y. co-conceived the study, designed, optimized and supervised the majority of experiments and analysis, performed oxidized DNA analysis in vitro and in cells, conducted cell death assays, drafted and revised the manuscript, and co-acquired funding. B.-X.Z. performed the majority of animal experiments, histological staining, and immunofluorescence; conducted all statistical analyses; co-analyzed data; prepared figures and contributed to manuscript drafting. H.-F.Y. maintained and expanded all mouse strains, performed a subset of mouse functional assays and tissue staining, co-performed Western blot analysis of tissue samples, and conducted quantitative real-time PCR. R.G. performed the majority of cell line-related experiments, including stimulation, Western blotting, subcellular fractionation, and mitochondrial DNA extraction, and co-performed tissue Western blot analysis. L.S. optimized mouse liver function assays and the APAP-induced injury model and co-acquired funding. Z.-Y.C. conducted EMSA experiments for oxidized DNA. Q.C. performed mitochondrial ROS detection, functional analysis, and flow cytometry. L.W. and J.H. generated gene-knockdown cell lines and rescue/overexpression cell lines. L.Z. constructed and prepared relevant plasmids. H.J. provided the MAVS knockout mouse strain. P.X. performed functional validation of cGAS knockout mice. Q.W. optimized dosing conditions for TH10785. J.Z. performed grayscale analysis of Western blots and quantitative fluorescence analysis. J.P. and S.F. co-performed mass spectrometry analysis of 8-oxodG and TH10785. H.Z. provided the *Ripk1*^+/−^ and *Fadd*^+/−^ mouse strains. X.S. assisted in project conceptualization, co-designed and co-conceived the study, and co-analyzed data. W.M. conceptualized the project, designed the study, analyzed data, reviewed and edited the manuscript, and acquired funding.

## Additional information

Correspondence and requests for materials should be addressed to Wei Mo (W.M.) or Zhang-Hua Yang (Z.-H.Y.)

---

**Extended Data Fig. 1.**
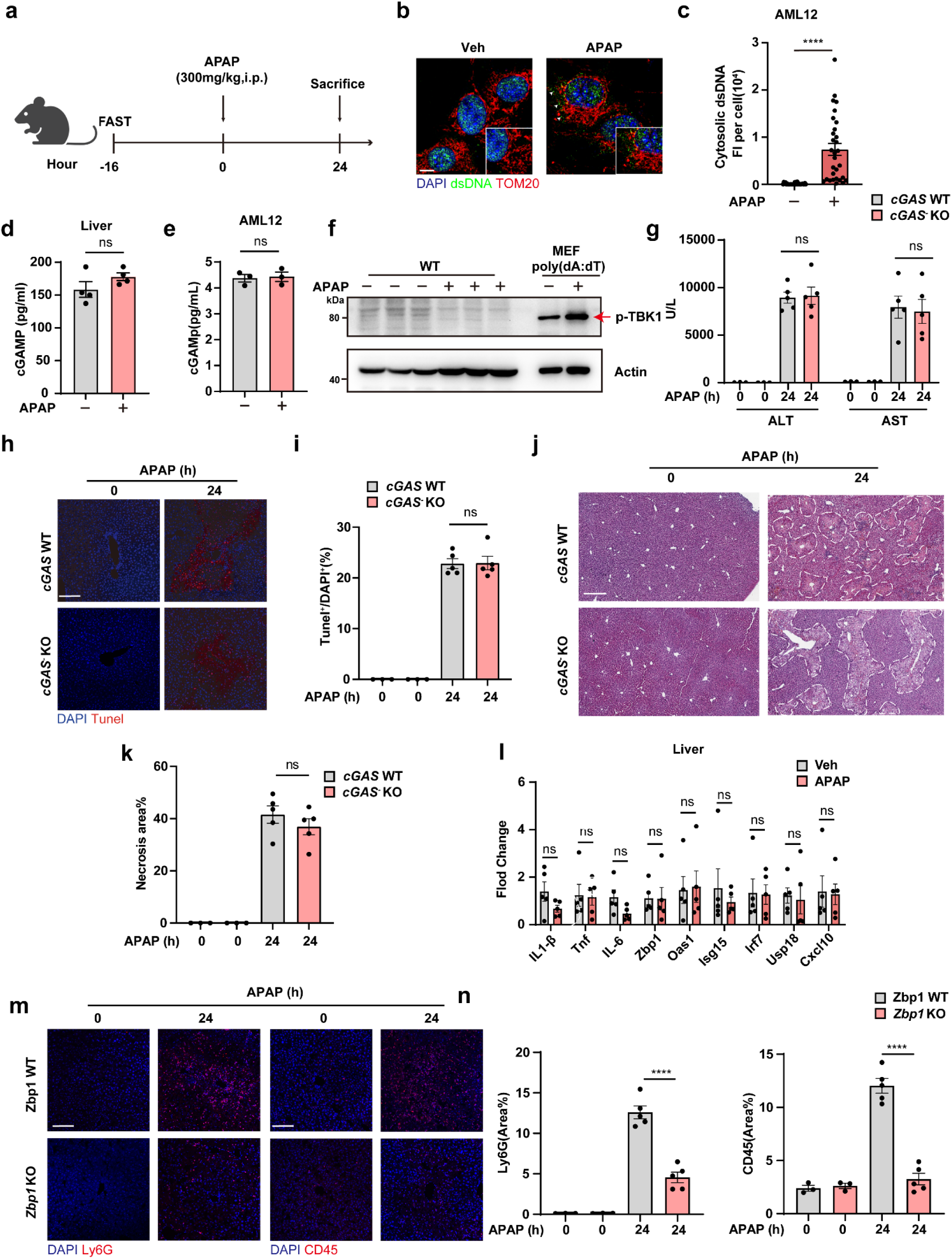
Cytoplasmic mtDNA activates ZBP1, not cGAS, to drive hepatocyte death. **a**, The scheme of APAP treatment. **b**, Immunofluorescence staining of dsDNA (green) and TOM20 (red) in AML12 cells treated with 10 mM APAP as indicated. The white arrow indicates the released mtDNA. Scale bar, 10 μm. **c**, Quantification of the fluorescence intensity of dsDNA in (**b**). n=10 cells per group, repeated 3 times independently with similar results. **d**, cGAMP production was measured using an enzyme-linked immunosorbent assay (ELISA) in liver of mice treated with APAP. n=4 mice per group. **e**, cGAMP production was measured using ELISA in AML12 treated with APAP. n=3 independent samples. **f**, Western blotting analysis of liver proteins from wildtype (WT) mice treated with or without APAP. n=3 mice per group. Protein sample from poly(dA:dT)-treated MEFs was used as the positive control. **g**, Serum ALT and AST levels were determined in *cGAS* WT and *cGAS* KO mice. 0 h: n=3 mice per genotype, 24 h: n=5 mice per genotype. **h**,**i**, Representative images (**h**) and quantification analyses (**i**) of TUNEL staining in liver tissue from *cGAS* WT and *cGAS* KO mice. 0 h: n=3 mice, 24 h: n = 5 mice. Scale bar, 100 μm. **j**, Hematoxylin and eosin (H&E) staining of the liver sections from *cGAS* WT and *cGAS* KO mice. **k**, Necrotic areas were encircled and quantified. 0 h: n=3 mice, 24 h: n=5 mice. Scale bar, 300 μm. **l**, qPCR analysis of ISGs, pro-inflammatory genes in the liver of WT mice treated with or without APAP. n=5 per group. **m**, Immunofluorescence staining for Ly6G and CD45 in *Zbp1* WT and *Zbp1* KO mice treated with APAP. Scale bar, 100 μm. **n**, Quantification analysis of the fluorescence area of Ly6G and CD45 in (**m**). 0 h: n=3 mice per genotype, 24 h: n=5 mice per genotype. Data are mean ± s.e.m. Statistical analysis was performed using Unpaired two-tailed Student’s *t*-test (**c, d** and **e**), two-way analysis of variance (ANOVA) (**g**; **i, k, l** and **n**), and one-way analysis of variance (ANOVA) (**i**); ns, not significant; \**P* < 0.05; \*\**P* < 0.01; \*\*\**P* < 0.001; \*\*\*\**P* < 0.0001.

**Extended Data Fig. 2.**
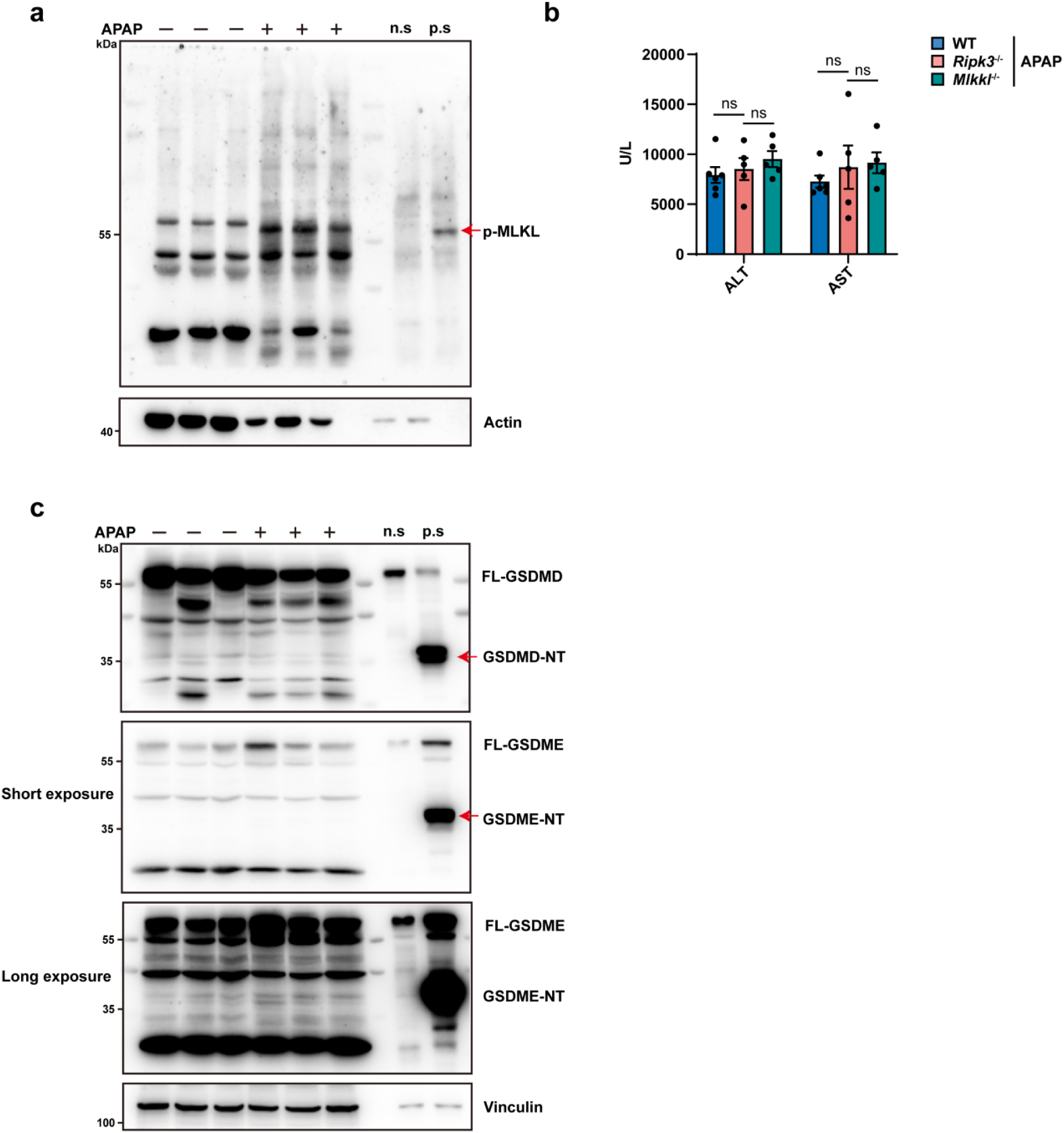
Neither necroptosis nor pyroptosis plays a role in APAP-induced hepatocyte death. **a**, Immunoblot analysis of liver lysates from WT mice treated as indicated for 24 h. Phosphorylated MLKL (p-MLKL) was detected in the RIPA insoluble fraction of liver lysates. Sample from TNFα+Smac+zVAD (TSZ)-treated MEFs was used as positive control for p-MLKL. n=3 mice per group. n.s: negative control; p.s: positive control. **b**, Serum ALT and AST levels were determined in WT, *Ripk3* KO, *Mlkl* KO mice treated with APAP. n=5 mice per group. **c**, Immunoblot analysis of liver lysates from WT mice treated as indicated for 24 h. n=3 mice per group. n.s: negative control; p.s: positive control. Protein sample from TNFα+Smac (TS)-treated MC38 cells was used as the positive control for GSDME-NT. Protein sample from LPS+Nigericin-treated RAW264.7 cells was used as the positive control for GSDMD-NT. Data are mean ± s.e.m. Statistical analysis was performed using two-way analysis of variance (ANOVA) (**b**), ns, not significant; \**P* < 0.05; \*\**P* < 0.01; \*\*\**P* < 0.001; \*\*\*\**P* < 0.0001.

**Extended Data Fig. 3.**
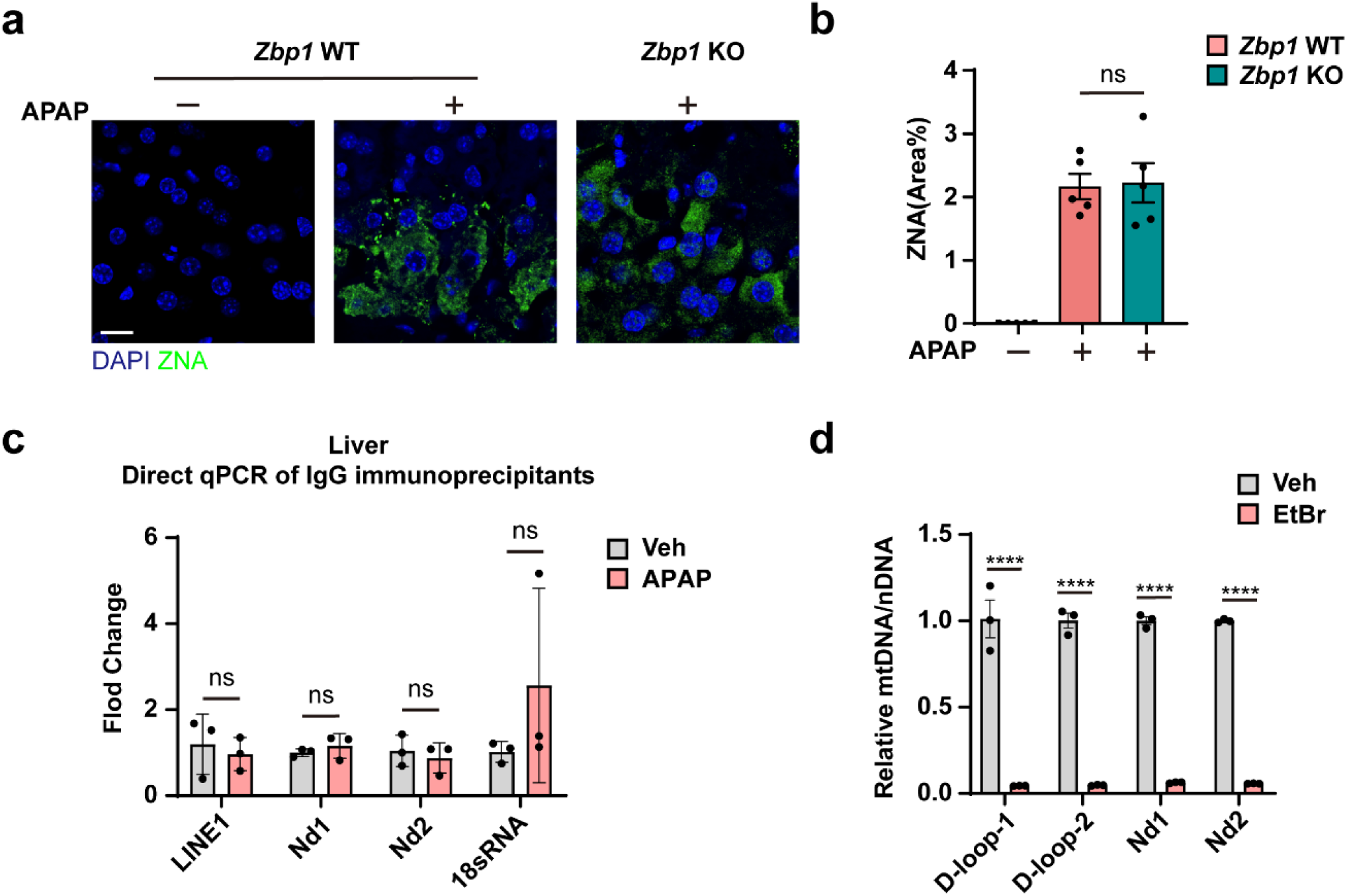
ZBP1 binds to Z-DNA derived from released mitochondrial DNA. **a**, Immunofluorescence staining of Z-NA (green) in the liver of *Zbp1* WT and *Zbp1* KO mice treated with or without APAP (300 mg/kg) for 24 h. Scale bars, 15 μm. **b**, Quantification of the fluorescence area of Z-NA signals in (**a**). n=5 mice per group. **c**, Quantitative qPCR analysis of ZBP1-binding DNAs enriched by anti-IgG immunoprecipitation in liver of WT mice treated with Vehicle or APAP (300 mg/kg) for 24 h. n=3 mice per group. **d**, Quantitative PCR analysis of cytosolic mtDNA release in cytosol fraction of AML12 cells with or without mtDNA depletion by EtBr (400 ng/ml) treatment for 6 days and stimulated with Vehicle or APAP (10 mM) for 24 h. Data are mean ± s.e.m. Statistical analysis was performed using one-way analysis of variance (ANOVA) (**b**), and two-way analysis of variance (ANOVA) (**c** and **d**); ns, not significant; \**P* < 0.05; \*\**P* < 0.01; \*\*\**P* < 0.001; \*\*\*\**P* < 0.0001.

**Extended Data Fig. 4.**
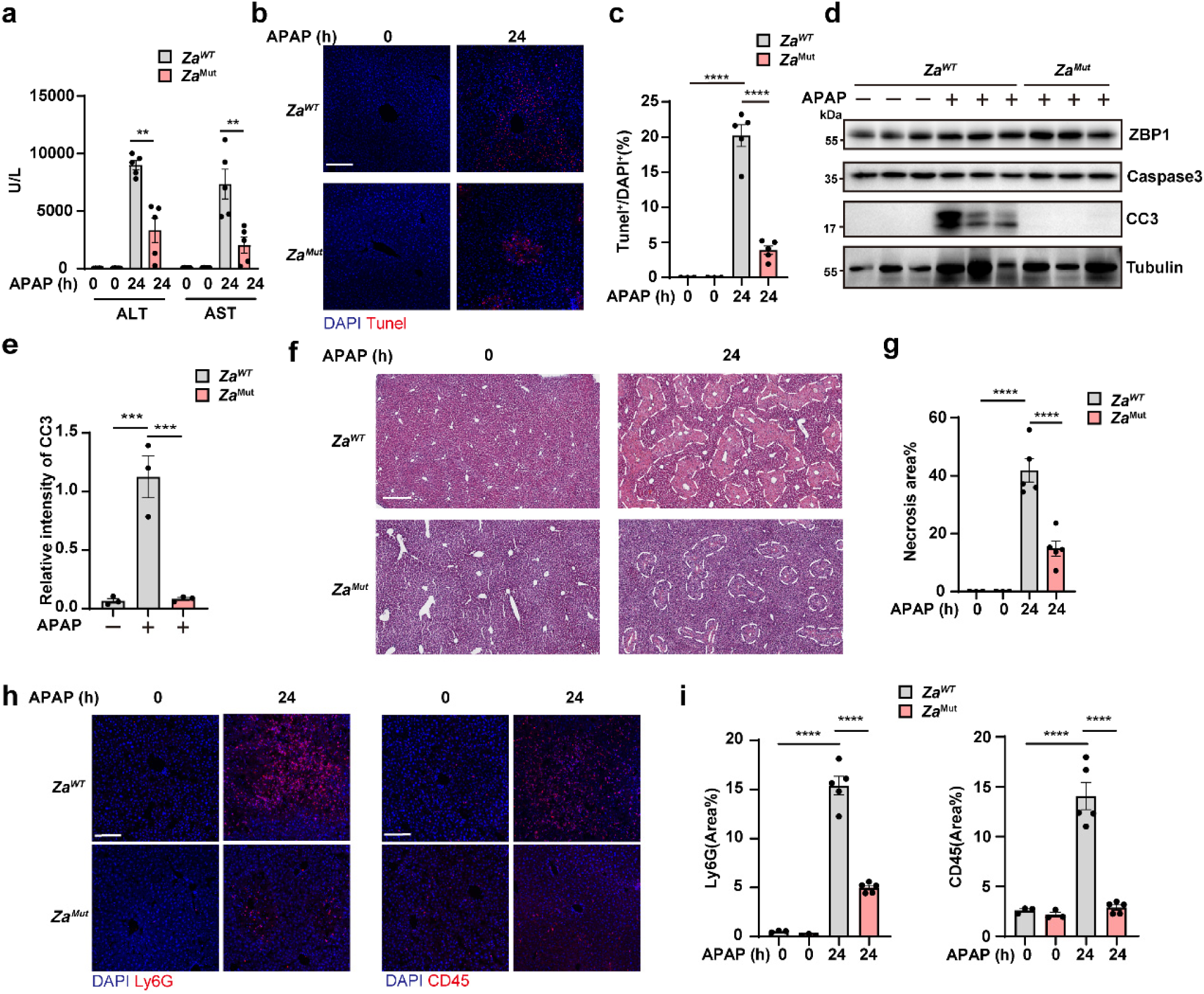
Z*α*-mediated recognition of Z-DNA by ZBP1 plays a pivotal role in APAP-induced hepatocyte death and liver injury. **a**, Serum ALT and AST levels were determined in *Za*^WT^ and *Za^Mut^* mice treated with APAP as indicated. 0 h: n=3 mice per genotype; 24 h: n=5 mice per genotype. **b**,**c**, Representative images (**b**) and quantification analyses (**c**) of TUNEL staining in liver tissue from *Za*^WT^ and *Za^Mut^* mice treated with APAP as indicated. 0 h: n=3 mice; 24 h: n = 5 mice. Scale bar, 100 μm. **d**,**e**, Western blotting (**d**) and quantification (**e**) analysis of liver proteins from *Za*^WT^ and *Za^Mut^* mice treated with APAP as indicated. n=3 mice per group. **f**, Hematoxylin and eosin (H&E) staining of the liver sections from *Za*^WT^ and *Za^Mut^*mice treated with APAP as indicated. **g**, Necrotic areas were encircled and quantified. 0 h: n=3 mice; 24 h: n=5 mice. Scale bar, 300 μm. **h**, Immunofluorescence staining for Ly6G and CD45 in *Zbp1* WT and *Zbp1* KO mice treated with APAP. Scale bar, 100 μm. **i**, Quantification analysis of the fluorescence area of Ly6G and CD45 in (**h**). 0 h: n=3 mice; 24 h: n=5 mice. Data are mean ± s.e.m. Statistical analysis was performed using one-way analysis of variance (ANOVA (**e**) and two-way analysis of variance (ANOVA) (**a, c, g** and **i**); ns, not significant; \*\**P* < 0.01; \*\*\**P* < 0.001; \*\*\*\**P* < 0.0001.

**Extended Data Fig. 5.**
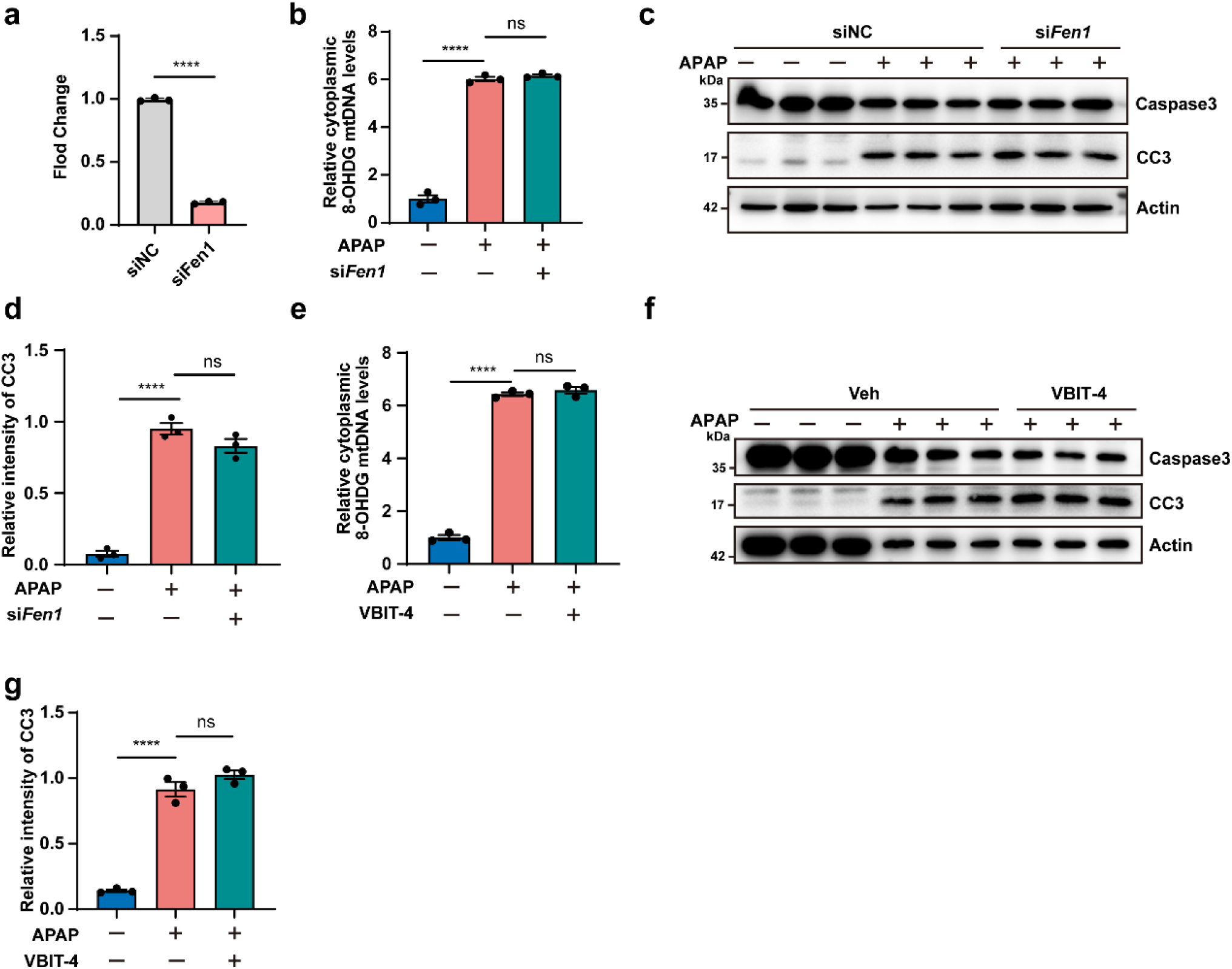
FEN1 and VDAC1 are dispensable for APAP-induced mtDNA fragmentation and cytoplasmic leakage. **a**, Quantitative RT-PCR analysis of *Fen1* in AML12 cells transfected with siRNAs targeting the indicated genes or non-target control (NC). n=3 independent samples. **b**, Quantification of the amounts of cytosolic Ox-mtDNA in the AML12 cells transfected with siRNAs targeting the indicated genes or NC and treated with or without APAP (10 mM) for 24 h. n=3 independent samples. **c**,**d**, Western blotting (**c**) and quantification (**d**) analysis of the AML12 cells transfected with siRNAs targeting the indicated genes or NC and treated with or without APAP (10 mM) for 24 h. n=3 independent samples. **e**, Quantification of the amounts of cytosolic Ox-mtDNA in the AML12 cells treated with or without APAP (10 mM) for 24 h, in the presence or absence of VBIT-4 (10 μM). n=3 independent samples. **f**,**g**, Western blotting (**f**) and quantification (**g**) analysis of the AML12 cells treated with or without APAP (10 mM) for 24 h, in the presence or absence of VBIT-4 (10 μM)). n=3 independent samples. Data are mean ± s.e.m. Statistical analysis was performed using unpaired two-tailed Student’s *t*-test (**a**) and one-way analysis of variance (ANOVA) (**b, d, e** and **g**); ns, not significant; \*\**P* < 0.01; \*\*\**P* < 0.001; \*\*\*\**P* < 0.0001.

**Extended Data Fig. 6.**
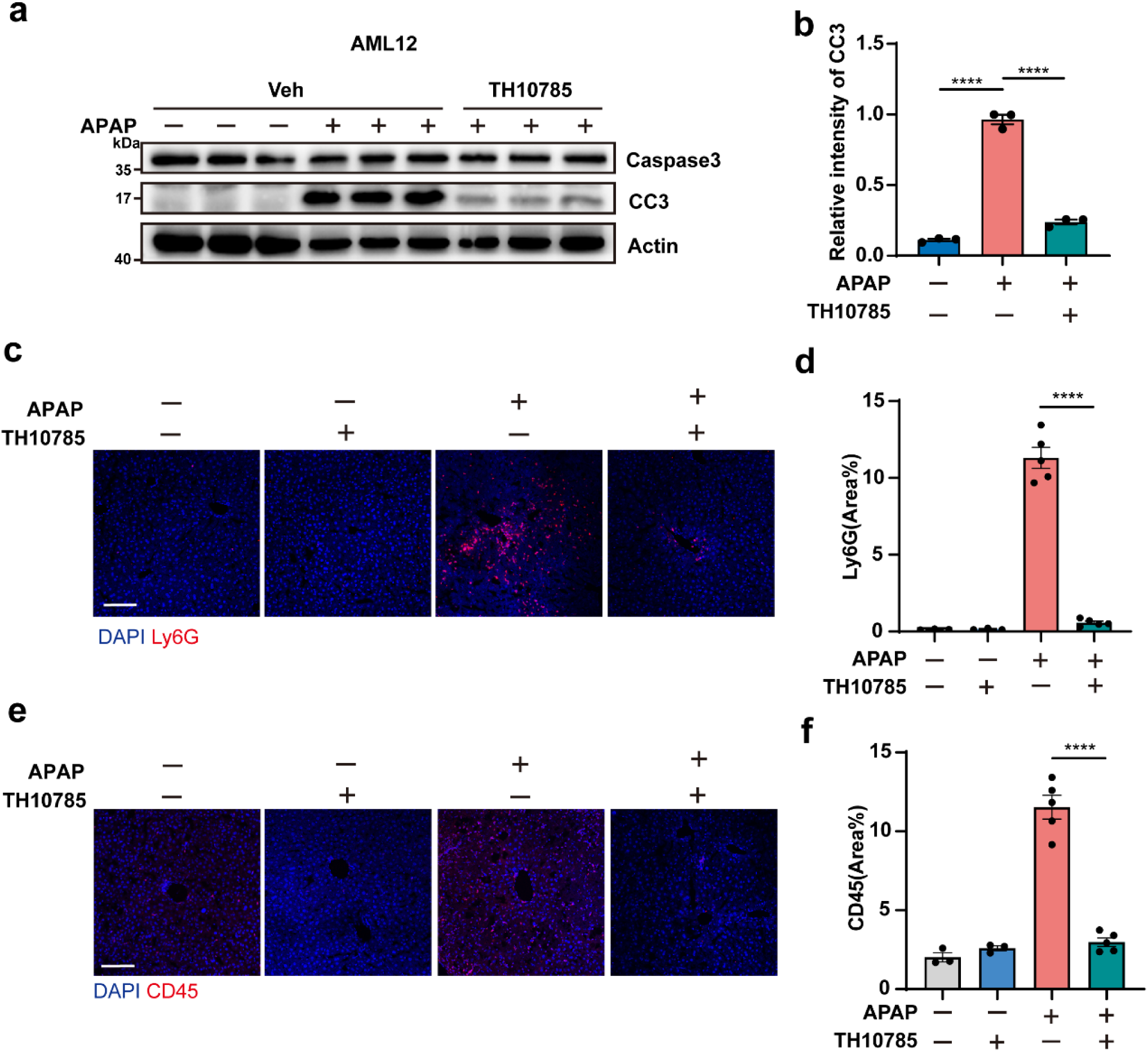
Deoxidation by OGG1 agonist TH10785 protects against APAP-induced cell death and liver failure. **a**,**b**, Western blotting (**a**) and quantification (**b**) analysis of the AML12 cells treated with or without APAP (10 mM) for 24 h, in the presence or absence of TH10785 (10 μM). n=3 independent samples. **c**,**d**, Immunofluorescence staining (**c**) and quantification (**d**) of Ly6G (red) in the liver of WT mice treated with or without APAP (300 mg/kg) for 24 h in the presence or absence of TH10785 (5 μg/g). Scale bars, 100 μm. **e**,**f**, Immunofluorescence staining (**e**) and quantification (**f**) of CD45 (red) in the liver of WT mice treated with or without APAP (300 mg/kg) for 24 h in the presence or absence of TH10785 (5 μg/g). **c-f**, WT mice only treated with or without TH10785: n=3; WT mice treated with or without APAP in the presence or absence of TH10785: n=5. Scale bars, 100 μm. Data are mean ± s.e.m. Statistical analysis was performed using one-way analysis of variance (ANOVA) (**b, d** and **f**); ns, not significant; \*\**P* < 0.01; \*\*\**P* < 0.001; \*\*\*\**P* < 0.0001.

**Extended Data Fig. 7.**
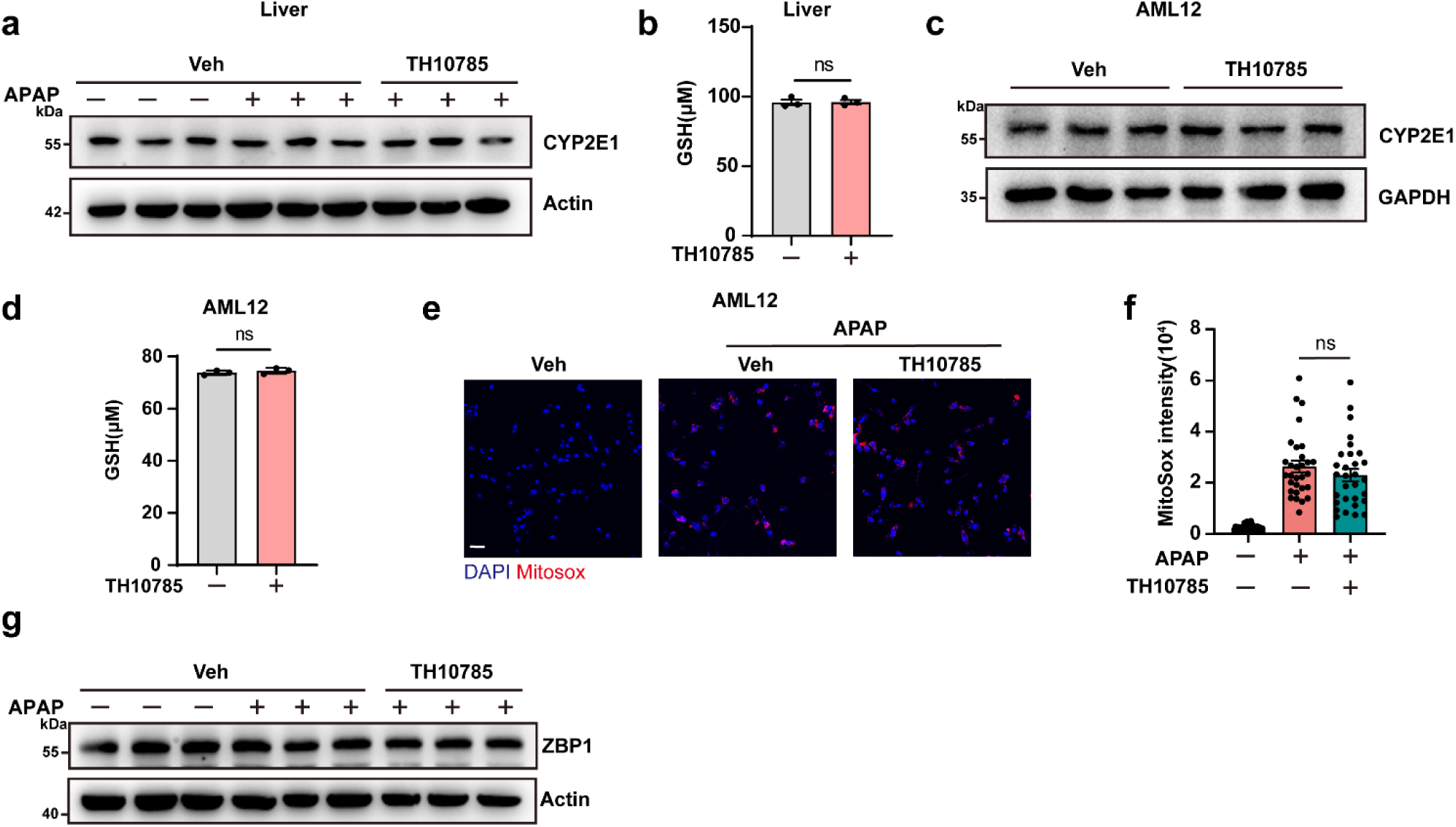
OGG1 agonist TH10785 did not affect APAP metabolism, ROS production or ZBP1 expression levels. **a**, Western blotting analysis of CYP2E1 in the liver from WT mice treated with or without APAP (300 mg/kg) for 24 h, in the presence or absence of TH10785 (10 μM). n=3 independent samples. **b**, Quantification analysis of GSH level in the liver from WT mice treated with or without TH10785 (10 μM) for 24 h. **c**, Western blotting analysis of CYP2E1 in AML12 cells treated with or without TH10785 (10 μM) for 24 h. n=3 independent samples. **d**, Quantification analysis of GSH level in AML12 cells treated with or without TH10785 (10 μM) for 24 h. n=3 independent samples. **e**, Representative images of mitoSOX Red+ cells indicating mitochondrial ROS production in AML12 cells treated with or without APAP (10 mM) for 24 h, in the presence or absence of TH10785 (10 μM). **f**, Quantification analysis of the fluorescence intensity of MitoSOX Red per cell in (**i**). n=10 cells per group, repeated 3 times independently with similar results. Scale bars, 50 μm. **g**, Western blotting analysis of ZBP1 in the AML12 cells treated with or without APAP for 24 h in the presence or absence of TH10785 (10 μM). n=3 independent samples. Data are mean ± s.e.m. Statistical analysis was performed using one-way analysis of variance (ANOVA) (**f**), and unpaired two-tailed Student’s *t*-test (**b** and **d**); ns, not significant; \*\**P* < 0.01; \*\*\**P* < 0.001; \*\*\*\**P* < 0.0001.

**Extended Data Fig. 8.**
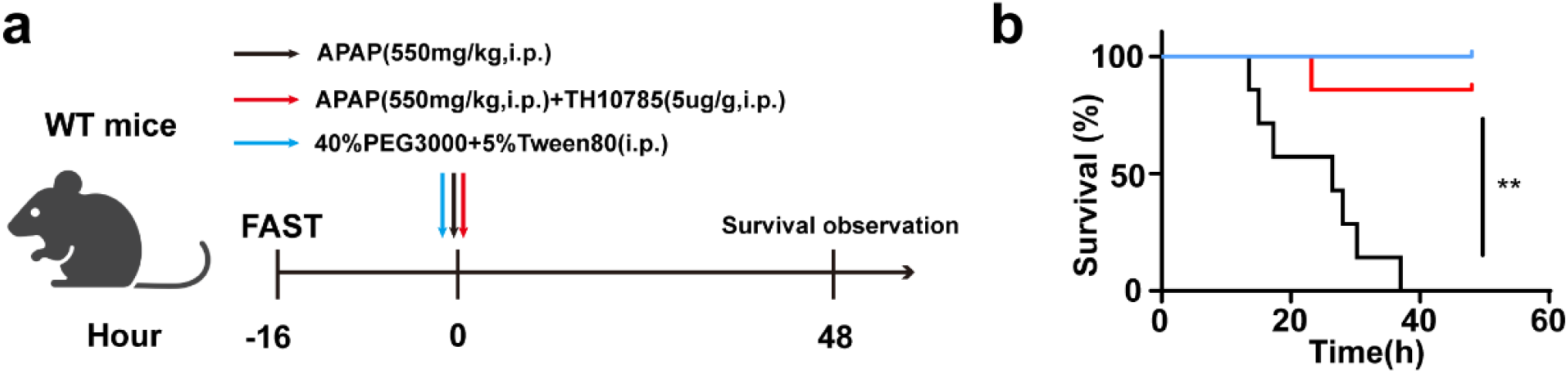
Deoxidation by OGG1 agonist TH10785 protects against APAP-induced mortality. **a**. The scheme of APAP treatment in the presence or absence of TH10785. **b**, Survival curves of WT mice treated with a lethal dose of APAP (550 mg/kg, i.p.) in the presence or absence of TH10785 (5 μg/g), n=7 mice per group. Data are mean ± s.e.m. Statistical analysis was performed using Mantel−Cox tests (**b**); ns, not significant; \*\**P* < 0.01; \*\*\**P* < 0.001; \*\*\*\**P* < 0.0001.

## Methods

### Animal experiments

All animal experiments were approved by the Animal Care and Use Committee of Zhejiang University. *Zbp1*^−/−^, *Ripk3*^−/−^, *Mlkl*^−/−^ mice were same as used^22^. *Zbp1^Za1,2Mut/Za1,2Mut^* mice were a gift from Dr. Jonathan. *cGAS^−/−^*mice were initially from The Jacksom Laboratory. C57BL/6 J mice (6- to 8-week-old) were purchased from from Shanghai SLAC Laboratory Animal Co., Ltd. All mice were on a C57BL/6 genetic background and housed in a constant temperature of 23℃ in a 12-hour light/dark cycle at the Laboratory Animal Center of Zhejiang University. They were fed with a standard laboratory diet (Research Diets, D12450K, USA) and had free access to water. To induce liver injury with APAP, male mice were used. They were fasted for 16 h and injected intraperitoneally (i.p.) with either saline or APAP (Yuanye Bio-Tech Co., Shanghai, China) at 300, or 550 mg/kg body weight. The animals were anesthetized via isoflurane induction at various timepoints after APAP dosing for analyses. To treat the animals with OGG1 agonist TH10785^4^, the compound was dissolved in DMSO (Sigma-Aldrich) at a concentration of 100 μg/uL as a stock solution and diluted with 40% PEG3000 and 5% Tween80 to 0.5 mg/ml before use. TH10785 was injected intraperitoneally at 5 μg/g. After the planned treatment, the mice were anesthetized, blood samples were collected from the retroorbital venous plexus for testing, and the livers were collected for analysis. The collected whole blood was centrifuged at 3000 g at 4°C for 10 min to obtain serum. The serum levels of alanine aminotransferase (ALT) and aspartate aminotransferase (AST) were determined with commercial kits (Sigma-Aldrich, Shanghai, China). The liver tissues were fixed in 4% paraformaldehyde, dehydrated and embedded in paraffin. Sections (7 µm) of the liver tissues were cut and collected. For histological examination, the sections were stained with hematoxylin and eosin (H&E). The necrotic areas in H&E-stained sections were encircled and measured with Adobe Photoshop under a 10x objective of each section were examined and measured to obtain the percentage of necrotic areas.

### Cell culture and stimulation

MEFs, AML12 cells were obtained in the lab. Cells were maintained in DMEM (Dulbecco’s modified Eagle’s medium) supplemented with 10% FBS (fetal bovine serum) and 1% (v/v) penicillin/streptomycin in an incubator supplied with a humidified atmosphere of 5% CO2 at 37 °C. For primary hepatocytes, the mouse liver was perfused with 0.05% collagenase type IV (Sigma-Aldrich) to obtain primary hepatocytes, which were plated in round coverslips in DMEM with 10% FBS and 1% PenStrep for 4 h for attachment. Subsequently, the cells were used in immunofluorescence staining. AML12 cells were incubated with APAP (10 mM) for 24 h to establish an in vitro DILI model. APAP was dissolved into serum free DMEM directly. Cells were seeded in six-well plates before being subjected to treatments. Supernatants and cell lysates were collected for ELISA and immunoblot (IB) analyses.

### Immunofluorescence staining

AML12 or MEF cells cultured in round coverslips were fixed in 4% PFA for 10 min and washed with PBS. After being permeabilized and blocked with 0.4% Triton X-100 and 3% BSA, cells were incubated with primary antibodies including 8oxodG (1:1000, Abcam, ab48508), ZNA (1:1000, Absolute Antibody, Ab00783-23.0), dsDNA (1:1000, Abcam, ab27156), Tom20 (1:1000, Proteintech, 11802-1-AP) antibody overnight at 4℃. Then the cells were washed with PBS, incubated with secondary antibody for 1 h at RT. Images were captured by Olympus FV3000.

### Tissue immunofluorescence

The liver tissues were fixed in 4% paraformaldehyde, dehydrated in 15% and 30% sucrose, and embedded in optimal cutting temperature compound at −20°C. The tissues were then sectioned (12 μm) using a cryostat. Afterwards, the sections were washed with PBS three times, blocked with 3% BSA and 0.4% Triton X-100 at RT for 1 h, and incubated with primary antibodies including CD45 (1:200, BD, 550539), Ly6G (1:200, Biolegend, 127601) overnight at 4℃. For ZNA staining in liver tissues, the tissues were pre-dried at 55℃ for 30 min and then subjected to proteinase K treatment (40 mg/mL) for 5 min at RT. When acquired, RNase A (5 mg/mL in PBS, Solarbio, 9001-99-4) or DNase I (25 U/mL in PBS, Invitrogen, AM2222) was used after proteinase K treatment at 37℃ for 1 h. The sections were incubated with primary antibodies (ZNA, 1:1000, Novus Biologicals, NB100-749) overnight at 4℃, washed with PBS three times, and incubated with secondary antibodies for 1 h at RT in the dark. Images were captured by Olympus FV3000 (fluorescence).

### TUNEL staining

The principle of TUNEL (TDT-mediated dUTP nick-end labelling) to detect cell apoptosis is that the exposed 3′-OH of broken DNA can be catalysed by terminal deoxynucleotidyl transferase (TdT) with FITC-labelled dUTP, which can be detected using fluorescence microscopy. The specific steps were performed according to the kit’s instructions (Vazyme, A113-03). Images were obtained by Olympus FV3000 (fluorescence). Digital images were recorded and analyzed using Image J software (NIH).

### Intensity line measurement

Representative images for intensity measurement were captured by Olympus FV3000, and pixel intensity was assessed with Fiji (ImageJ) to measure indicated staining (ZNA/8oxodG or ZNA/8oxodG/FAM) intensity across the dotted lines at baseline and after APAP-treated AML12 cells or FAM-8oxo-dG_3_dC duplexes-transfected cells.

### Automatic image analysis

For the quantification of mitochondrial superoxide, the intensity of the mitochondrial superoxide indicator MitoSOX was analyzed within cells using Hoechst as the nuclear counterstain and defining cells by propagation of the nuclear objects to the cellular periphery on the basis of MitoSOX staining. For the quantification of cytoplasmic dsDNA, the intensity of the cytoplasmic dsDNA fluorescence intensity was analyzed within cells using Hoechst as the nuclear counterstain and defining cells by propagation of the nuclear objects to the cellular periphery on the basis of dsDNA staining. The statistical analysis of ZNA, CD45, Ly6G fluorescence is performed by using ImageJ to calculate the percentage of ZNA, CD45, Ly6G fluorescence in the field of view. To quantify the colocalization of ZNA/8oxodG, FAM/ZNA or FAM/8oxodG, the Manders’ colocalization coefficient (MCC) was calculated using ImageJ software. First, dual-channel fluorescence images under a 60x objective were acquired under identical acquisition settings to ensure consistency. The images were then opened in ImageJ, and background subtraction was performed to minimize noise interference. The “Coloc JACop” plugin in ImageJ was utilized for the analysis.

### Measurement of B-DNA and Z-DNA conformation using a ratio of A260/295

Conformation of Z-DNA and B-DNA was assessed using the absorbance ratio of 260 to 295 nm as previously described ^28,29^. We used short double-stranded DNA (dsDNA) sequences (12 bp) with six consecutive GC repeats with defined oxidation levels (0, 1, or 3 8-oxodG substitutions per strand) (100 ng/μL) for conformation analysis. Poly(dG:dC) was diluted in annealing buffer (20 mM NaCl, 20mM Tris-HCL, 0.1mM EDTA) to induce Z-DNA conformation. A260/295 was measured with a NanoDrop (Thermo Fisher Scientific) using annealing buffer as a blank. Each incubation was performed in triplicates. The values were then plotted as the ratio of A260/295.

### dsDNA transfection

For dsDNA transfection in 96-well plates, 0.1 μg of DNA duplexes were mixed with Lipofectamine 2000 (Thermo, 11668030) and brought up to 20 μL in OptiMEM and allowed to sit at room temperature for 20 min. The mixture was then added to the MEF cells and allowed to incubate at 37℃ for 8 h. The medium was changed 6 h post transfection.

### Generation of mitochondrial DNA-depleted AML12 cells

To generate mitochondrial DNA-depleted AML12, AML12 were cultured in medium containing 400 ng/ml of ethidium bromide (EtBr, Sangon, A500328) for 6 days prior to the indicated treatments. To measure the efficiency of mtDNA depletion, total DNA were extracted for real-time qPCR analysis to measure the expression of mitochondrial genes (*D-loop-1*, *D-loop-2*, *Nd1*, and *Nd2*), and the expression values of each replicate were normalized against nuclear-encoded *Tert*.

### Electrophoretic Mobility Shift Assay (EMSA)

The binding assays were performed by incubating 2.5 µM of FAM-dGdC, FAM-8oxo-dG_1_dC, and FAM-8oxo-dG_3_dC oligonucleotides in 10 mM HEPES buffer (pH 7.5, containing 10 mM MgCl_2_) with either reconstituted Human-ZBP1-Zα1 or its mutant form (Human-ZBP1-Zα1-mutant) at specified DNA:protein molar ratios. The incubation was carried out for 2.5 hours at 25°C. Following incubation, the DNA-protein complexes were separated by native polyacrylamide gel electrophoresis using 15% TBE gels. DNA visualization was achieved through post-staining with ethidium bromide, and the resulting bands were captured using the Magic SHST‘s Gel doc system.

### *Zbp1* knockdown by shRNA lentivirus

Sequences of specific shRNAs used in this study were obtained from the MISSION shRNA Library (Sigma). Lentiviral particles generated by co-transfection psPAX2 and PMD2.G packing plasmids into HEK293T cells were used to knockdown *Zbp1* in AML12 cells. Supernatants were collected 48 h after transfection, filtrated through a 0.45 µm pore filter and added to AML12 cells. To increase infection efficiency, 8 μg/mL of polybrene were added. The virus containing medium was washed after 12 h and the cells were cultured with fresh medium. Infected cells were expanded and selected with puromycin at 48 h post-transduction.

### siRNA transfection in AML12

For siRNA transfection in 24-well plates, a mixture containing 0.75 μL of siRNA duplexes (20 μM), 1.5 μL of Lipofectamin RNAiMAX reagent (Thermo, 13778150), and 75 μL of Opti-MEM (Gibco, 31985070) was incubated for 5 min and then added into AML12 culture for 24 h before treatment. The following oligonucleotide sequences were used for knockdown: si*Fen1*, 5’-CGUGCUAAUGCGACACUUATT-3’; siNC, 5’-UUCUCCGAACGUGUCACGUTT-3’.

### Immunoblotting

For mouse liver tissues, samples were collected from mice following cardiac perfusion with PBS, tissues were homogenized using metal beads at 60 Hz for 60 s in ice-cold RIPA lysis buffer (50 mM Tris-HCl pH 7.5, 150 mM NaCl, 1% NP-40, 0.5% sodium deoxycholate, 0.1% SDS, 1% protease inhibitor cocktail, 1 mM PMSF, and 1% phosphatase inhibitor) with a TissueLyser II (QIAGEN), and centrifuged at 12,000 ×g at 4℃ for 30 min. Then the protein concentrations were adjusted to 1 mg/ml based on BCA. Proteins were blotted following a standard protocol. Antibodies against the following proteins were used for immunoblotting: ZBP1 (AdipoGen, AG-20B-0010-C100, 1:1000), p-TBK1 S172 (CST, 5483S, 1:1000), Cleaved Caspase-3 (CST, 9664S, 1:1000), Caspase-3 (CST, 9662S, 1:1000), p-MLKL (phospho S345)(Abcam, ab196436, 1:1000), MLKL (Proteintech, 66675-1-lg, 1:1000), GSDME (Abcam, ab215191,1:1000), GSDMD (Abcam, ab209845, 1:1000), Actin (Proteintech, 66009-1-Ig, 1:5000), GAPDH (HUABIO, ET1601-4, 1:5000), Vinculin (HUABIO, ET1705-94, 1:5000) and Tubulin (Millipore, 05-829, 1:10000). The signals were detected by Immobilon ECL Ultra Western HRP Substrate (Millipore).

### ELISA

For the measurement of cGAMP, AML12 cells were digested and collected into centrifuge tubes, and cells were counted using the Countstar automated after two washes with cold PBS. Cells were lysed by repeated freeze-thaw cycles in PBS at a ratio of 100 μL per million cells. Mice were euthanized, and the liver tissues were carefully dissected. A total of 1mL of PBS was used for every 100 mg of fresh mouse liver tissue, which were then thoroughly disrupted using Tissuelyser-II (Shanghai Jingxin). Tissue homogenates were placed into a 3D shaker at 4°C for 5 min before centrifuging. The Pierce BCA Protein Assay was used to normalize the protein concentration of the supernatants. The 2′3′-cGAMP concentrations were measured using ELISA (COIBO, CB15107-Mu) according to the manufacturer’s instructions. For the measurement of Ox-mtDNA, purified mtDNA was extracted from the cytosolic or mitochondrial fractions as indicated. The 8-OH-dG content was then quantified using 8-hydroxy 2-deoxyguanosine ELISA Kit (COIBO BIO, CB10013) according to manufacturer’s instructions.

### Cytosolic DNA immunoprecipitation

Liver tissues were lysed at 4℃ for 10 min with 1 mL of digitonin lysis buffer (150 mM NaCl, 20 mM HEPES pH7.4, and 25 μg/mL digitonin). Tissue lysates were centrifuged twice at 13,000 × g for 5 minutes at 4℃ to separate the soluble fraction from a pellet that contains the heavy membrane fraction. Immunoprecipitation of ZBP1 was performed using anti-ZBP1 and IgG. Samples were incubated at 4℃ overnight on a rotator, washed extensively and then subjected to phenol-chloroform based DNA extraction. Quantification of co-precipitated DNA was analyzed by quantitative real-time qPCR, and presented as the relative expression of untreated control.

### Quantitative real-time PCR

Total RNA was isolated from mice liver using TRIzol reagent (Life Technologies). RNA concentration was measured using the Nanodrop spectrophotometer (Thermo). cDNA was prepared using 1 μg of RNA with HiScript III room temperature SuperMix kit (Vazyme), and reverse transcribed into cDNA. qPCR was performed with ChamQ Universal SYBR qPCR Master Mix (Vazyme) by the CFX Connect Real-Time PCR Detection System (Bio-Rad). Data were analyzed according to the ΔΔ*CT* method. *Actin* (encoding β-actin) was used as the reference gene for accurate normalization of qPCR data. To measure the abundance of cytosolic mtDNA, 10 ng/μL of template DNA was used for qPCR analysis, and expression values of each replicate (*D-loop-1*, *D-loop-2*, *Nd1*, and *Nd2*) were normalized against nuclear-encoded *Tert*. The sequences of gene-specific primers used for PCR are shown below.

*Zbp1*-F:5’-TTGAGCACAGGAGACAATCTG-3’
*Zbp1*-R:5’-TTCAGGCGGTAAAGGACTTG-3’;
*Ifnb1*-F: 5’-CAGCTCCAAGAAAGGACGAAC-3’,
*Ifnb1*-R: 5’-GGCAGTGTAACTCTTCTGCAT-3’;
*Tnf*-F: 5’-CCCTCACACTCAGATCATCTTCT-3’,
*Tnf*-R: 5’-GCTACGACGTGGGCTACAG-3’;
*Il6*-F: 5’-TAGTCCTTCCTACCCCAATTTCC-3’,
*Il6*-R: 5’-TTGGTCCTTAGCCACTCCTTC-3’;
*Il1b*-F: 5’-GCAACTGTTCCTGAACTCAACT-3’,
*Il1b*-R: 5’-ATCTTTTGGGGTCCGTCAACT-3’;
*Usp18*-F: 5’-TTGGGCTCCTGAGGAAACC-3’,
*Usp18*-R: 5’-CGATGTTGTGTAAACCAACCAGA-3’;
*Isg15*-F: 5’-CCCCCATCATCTTTTATAACCAAC-3’,
*Isg15*-R:5’-CACAGTGATCAAGCATTTGCG-3’;
*Oasl1*-F: 5’-TCCTTCGGTTGGTCAAACAC-3’,
*Oasl1*-R: 5’-CAGGCATAGACAGTGAGCAG-3’;
*Irf7*-F: 5’-TGTTTGGAGACTGGCTATTGG-3’,
*Irf7*-R:5’-ATCCCTACGACCGAAATGCT-3’;
*Tert*-F: 5’-CTAGCTCATGTGTCAAGACCCTCTT-3’,
*Tert*-R:5’-GCCAGCACGTTTCTCTCGTT-3’;
*D-loop-1-*F: 5’-AATCTACCATCCTCCGTGAAACC-3’,
*D-loop-1-*R: 5’-TCAGTTTAGCTACCCCCAAGTTTAA-3’;
*D-loop-2-*F: 5’-TCCTCCGTGAAACCAACAA-3’,
*D-loop-2-*R: 5’-AGCGAGAAGAGGGGCATT-3’;
*Nd1*-F: 5’-CTAGCAGAAACAAACCGGGC-3’,
*Nd1*-R: 5’-CCGGCTGCGTATTCTACGTT-3’;
*Nd2*-F: 5’-CCATCAACTCAATCTCACTTCTATG-3’,
*Nd2*-R: 5’-GAATCCTGTTAGTGGTGGAAGG-3’;
*Fen1*-F: 5’-TTCACGGCCTTGCCAAACTAA-3’,
*Fen1*-R: 5’-ACAGCAATCAGGAACTGGTAGA-3’;
*LINE1*-F: TAGGAAATTAGTTTGAATAGGTGAGAGGGT,
*LINE1*-R: TCCAGAAGCTGTCAGGTTCTCTGGC;
*RNA18S*-F: GTAACCCGTTGAACCCCATT,
*RNA18S*-R: CCATCCAATCGGTAGTAGCG.
*Actin*-F: 5’-GGCTGTATTCCCCTCCATCG-3’,
*Actin*-R: 5’-CCAGTTGGTAACAATGCCATGT-3’;

### Measurement of mitoROS

MitoSOX™ (Invitrogen, USA) red dye is living-cell permeant and is capable of selectively targeting mitochondria where once it was oxidized by superoxide, it would produce red fluorescence (λex = 396 nm; λem = 610 nm). AML12 cells were cultured in 35mm dish, the medium was discarded after treated with 10 mM APAP for 24 h, and the cells were washed with PBS for 3 times. 1 mL of 5 μM MitoSOX solution was added to cells and incubate at 37°C for 30 min. Then, cells were washed again with PBS for 3 times, and dyed with 1 μg/mL Hoechst for 10 min. Images were captured by Olympus FV3000 (fluorescence).

### Measurement of GSH

To examine GSH, mouse liver tissue was homogenized with extraction solution at the following ratio: weight (g):volume (mL) = 1:10. The supernatant was collected by centrifugation at 10,000g at 4°C for 10 min. Protein concentration was determined by a BCA detection kit, and the levels of GSH in liver tissues were detected by using a commercial GSH kit (S0053, Beyotime Biotechnology) in accordance with the instructions. Cells were seeded in 6-well plates and cultured overnight. After indicated treatments, cells were harvested and cell numbers were determined. The identical number of cells (2 × 106 cells for each sample) were taken for further GSH measurement by using a commercial GSH kit (S0053, Beyotime Biotechnology).

### Quantification and statistical analysis

Data for cell experiments are presented as mean ± SEM of the indicated number of independent samples in one representative experiment. Mouse data are presented as mean ± SEM of the indicated *n* values. Statistical parameters including the exact sample size (*n*), post hoc tests, and statistical significance are reported in every figure and figure legend. Statistical analyses were performed with GraphPad Prism 8.0 using two-tailed unpaired Student’s t-tests, one-way ANOVA or two-way ANOVA, as indicated in the figure legends. The Mantel–Cox test was used for comparing mouse survival curves. Differences were considered statistically significant if *P* < 0.05.

## References

1 Smilkstein, M. J., Knapp, G. L., Kulig, K. W. & Rumack, B. H. Efficacy of oral N-acetylcysteine in the treatment of acetaminophen overdose. Analysis of the national multicenter study (1976 to 1985). N Engl J Med 319, 1557–1562 (1988). 10.1056/NEJM198812153192401

2 Yoon, E., Babar, A., Choudhary, M., Kutner, M. & Pyrsopoulos, N. Acetaminophen-Induced Hepatotoxicity: a Comprehensive Update. J Clin Transl Hepatol 4, 131–142 (2016). 10.14218/JCTH.2015.00052

3 Fromme, J. C., Bruner, S. D., Yang, W., Karplus, M. & Verdine, G. L. Product-assisted catalysis in base-excision DNA repair. Nat Struct Biol 10, 204–211 (2003). 10.1038/nsb902

4 Michel, M. et al. Small-molecule activation of OGG1 increases oxidative DNA damage repair by gaining a new function. Science 376, 1471–1476 (2022). 10.1126/science.abf8980

5 Tran, T. & Lee, W. M. DILI: New Insights into Diagnosis and Management. Curr Hepat Rep 12, 53–58 (2013). 10.1007/s11901-012-0159-x

6 Han, D. et al. Regulation of drug-induced liver injury by signal transduction pathways: critical role of mitochondria. Trends Pharmacol Sci 34, 243–253 (2013). 10.1016/j.tips.2013.01.009

7 Nelson, S. D. Molecular mechanisms of the hepatotoxicity caused by acetaminophen. Semin Liver Dis 10, 267–278 (1990). 10.1055/s-2008-1040482

8 Kon, K., Kim, J. S., Jaeschke, H. & Lemasters, J. J. Mitochondrial permeability transition in acetaminophen-induced necrosis and apoptosis of cultured mouse hepatocytes. Hepatology 40, 1170–1179 (2004). 10.1002/hep.20437

9 Umbaugh, D. S., Nguyen, N. T., Jaeschke, H. & Ramachandran, A. Mitochondrial Membrane Potential Drives Early Change in Mitochondrial Morphology After Acetaminophen Exposure. Toxicol Sci 180, 186–195 (2021). 10.1093/toxsci/kfaa188

10 McGill, M. R. & Jaeschke, H. Metabolism and disposition of acetaminophen: recent advances in relation to hepatotoxicity and diagnosis. Pharm Res 30, 2174–2187 (2013). 10.1007/s11095-013-1007-6

11 Newman, L. E. & Shadel, G. S. Mitochondrial DNA Release in Innate Immune Signaling. Annu Rev Biochem 92, 299–332 (2023). 10.1146/annurev-biochem-032620-104401

12 Hopfner, K. P. & Hornung, V. Molecular mechanisms and cellular functions of cGAS-STING signalling. Nat Rev Mol Cell Biol 21, 501–521 (2020). 10.1038/s41580-020-0244-x

13 Lauterbach-Riviere, L. et al. Hepatitis B Virus DNA is a Substrate for the cGAS/STING Pathway but is not Sensed in Infected Hepatocytes. Viruses 12 (2020). 10.3390/v12060592

14 Thomsen, M. K. et al. Lack of immunological DNA sensing in hepatocytes facilitates hepatitis B virus infection. Hepatology 64, 746–759 (2016). 10.1002/hep.28685

15 Choubey, D. Cytosolic DNA sensor IFI16 proteins: Potential molecular integrators of interactions among the aging hallmarks. Ageing Res Rev 82, 101765 (2022). 10.1016/j.arr.2022.101765

16 Lugrin, J. & Martinon, F. The AIM2 inflammasome: Sensor of pathogens and cellular perturbations. Immunol Rev 281, 99–114 (2018). 10.1111/imr.12618

17 Zhang, Z. et al. The helicase DDX41 senses intracellular DNA mediated by the adaptor STING in dendritic cells. Nat Immunol 12, 959–965 (2011). 10.1038/ni.2091

18 Ishii, K. J. et al. TANK-binding kinase-1 delineates innate and adaptive immune responses to DNA vaccines. Nature 451, 725–729 (2008). 10.1038/nature06537

19 Amusan, O. T. et al. RIPK1 is required for ZBP1-driven necroptosis in human cells. PLoS Biol 23, e3002845 (2025). 10.1371/journal.pbio.3002845

20 Yang, D. et al. ZBP1 mediates interferon-induced necroptosis. Cell Mol Immunol 17, 356–368 (2020). 10.1038/s41423-019-0237-x

21 Zhang, T. et al. Influenza Virus Z-RNAs Induce ZBP1-Mediated Necroptosis. Cell 180, 1115–1129 e1113 (2020). 10.1016/j.cell.2020.02.050

22 Wang, R. et al. Gut stem cell necroptosis by genome instability triggers bowel inflammation. Nature 580, 386–390 (2020). 10.1038/s41586-020-2127-x

23 Lei, Y. et al. Cooperative sensing of mitochondrial DNA by ZBP1 and cGAS promotes cardiotoxicity. Cell 186, 3013–3032 e3022 (2023). 10.1016/j.cell.2023.05.039

24 Jiao, H. et al. Z-nucleic-acid sensing triggers ZBP1-dependent necroptosis and inflammation. Nature 580, 391–395 (2020). 10.1038/s41586-020-2129-8

25 Xian, H. et al. Oxidized DNA fragments exit mitochondria via mPTP- and VDAC-dependent channels to activate NLRP3 inflammasome and interferon signaling. Immunity 55, 1370–1385 e1378 (2022). 10.1016/j.immuni.2022.06.007

26 Kim, J. et al. VDAC oligomers form mitochondrial pores to release mtDNA fragments and promote lupus-like disease. Science 366, 1531–1536 (2019). 10.1126/science.aav4011

27 Zhang, T. et al. ADAR1 masks the cancer immunotherapeutic promise of ZBP1-driven necroptosis. Nature 606, 594–602 (2022). 10.1038/s41586-022-04753-7

28 Chaires, J. B. Allosteric conversion of Z DNA to an intercalated right-handed conformation by daunomycin. J Biol Chem 261, 8899–8907 (1986).

29 Buzzo, J. R. et al. Z-form extracellular DNA is a structural component of the bacterial biofilm matrix. Cell 184, 5740–5758 e5717 (2021). 10.1016/j.cell.2021.10.010

